# Identifying transcription factor-bound activators and silencers in the chromatin accessible human genome using ATAC-STARR-seq

**DOI:** 10.1101/2022.03.25.485870

**Authors:** Tyler J. Hansen, Emily Hodges

## Abstract

Massively parallel reporter assays test the capacity of putative gene regulatory elements to drive transcription on a genome-wide scale. Most gene regulatory activity occurs within accessible chromatin, and recently described methods have combined assays that capture these regions—such as assay for transposase-accessible chromatin using sequencing (ATAC-seq)—with self-transcribing active regulatory region sequencing (STARR-seq) to selectively assay the regulatory potential of accessible DNA (ATAC-STARR-seq). Here, we report a multi-omic approach that quantifies regulatory activity, chromatin accessibility, and transcription factor (TF) occupancy with one assay using ATAC-STARR-seq. Our strategy, including important updates to the ATAC-STARR-seq assay design and workflow, enabled high-resolution testing of ∼50 million unique DNA fragments tiling ∼101,000 accessible chromatin regions in human lymphoblastoid cells. We discovered that 30% of all accessible regions contain an activator, a silencer or both. We demonstrate that activators and silencers represent distinct functional groups that are enriched for unique sets of TF motifs and are marked by specific combinations of histone modifications. Using Tn5 cut-sites retained by the ATAC-STARR library, we performed TF footprinting and stratified these groups by the presence of specific TF footprints that are supported by chromatin immunoprecipitation data. We found that activators and silencers clustered by distinct TF footprint combinations are enriched for distinct gene regulatory pathways, and thus, represent distinct gene regulatory networks of human lymphoblastoid cell function. Altogether, these data highlight the multi-faceted capabilities of ATAC-STARR-seq to comprehensively investigate the regulatory landscape of the human genome all from a single DNA fragment source.

## INTRODUCTION

Transcription is regulated by transcription factors (TFs) and the DNA sequences they bind, called *cis*-regulatory elements. To regulate transcription, TFs bind *cis*-regulatory elements at their cognate sequence motifs and interact with co-activating factors to promote both the assembly of the pre-initiation complex and its release into productive elongation (Haberle and Stark 2018; Lambert et al. 2018). Enhancers are *cis*-regulatory elements distally located from the genes they target and serve as key drivers of cell-type specific gene expression (Heinz et al. 2015). Because enhancers require TF binding, they are largely dependent on chromatin accessibility to elicit transcriptional activity. Therefore, chromatin accessibility is a vital regulator of enhancer function, and this is evidenced by the presence of ∼94% and ∼97% of all ENCODE TF ChIP-seq peaks and distal *cis*-regulatory elements compiled from 1,046 ENCODE datasets within accessible chromatin, respectively (Thurman et al. 2012; Klemm et al. 2019). In any given cell type, only a small fraction (∼2%) of the genome is accessible to TF binding (Thurman et al. 2012; Klemm et al. 2019). In this way, most enhancers are inaccessible and are less likely to drive transcription endogenously. Critically, however, chromatin accessibility alone is not sufficient to drive or repress transcriptional activity and enhancers are dependent on additional regulatory processes that control TF binding (Dogan et al. 2015; Consortium et al. 2020).

Unlike their promoter counterparts, which are located proximal to transcription start sites, enhancers are difficult to identify and validate because they lack uniform features and are less constrained by gene proximity (Gasperini et al. 2020). The most robust way to test enhancer function, which is the ability of the enhancer to drive transcription of a target gene, is to place enhancer sequences in their native context in a transgenic animal model; however, it is fundamentally low throughput to do so. Instead, cell-based plasmid reporter assays offer a higher-throughput approach to directly test and validate a putative enhancer sequence. However, these assays cannot accommodate or functionally dissect the activity of millions of putative enhancers (Ryan and Farley 2020). To increase throughput, massively parallel reporter assays (MPRAs) were developed to test the regulatory potential of thousands to millions of DNA sequences in parallel by cloning them *en masse* into a reporter plasmid and leveraging next-generation sequencing to quantify regulatory activity (Santiago-Algarra et al. 2017). Among the variety of different vector backbones and assay designs applied to MPRAs, Self-Transcribing Active Regulatory Region sequencing (STARR-seq) is uniquely designed to assay an entire genome for regulatory activity (Melnikov et al. 2012; Patwardhan et al. 2012; Arnold et al. 2013; Inoue et al. 2017; Maricque et al. 2017; Muerdter et al. 2018; Kircher et al. 2019).

STARR-seq quantifies regulatory activity genome-wide by cloning randomly fragmented genomic DNA into the 3’UTR of the reporter plasmid. Thus, active enhancers drive transcription of themselves, and activity is quantified by the abundance of its own sequence in the transcript pool, removing the need for barcodes that some MPRAs employ. One major limitation of STARR-seq is that it is technically challenging to accommodate the massive size of the human genome; it requires large-scale cloning procedures and produces shallow sequencing coverage of human regulatory elements (Johnson et al. 2018). In addition, STARR-seq assays both accessible and inaccessible chromatin. Thus, many assayed regions are derived from heterochromatin and are less likely to be transcriptionally active in the cell type in question.

To narrow the scope of STARR-seq while assaying nearly all endogenously active regulatory elements, recent methods, such as assay for transposase-accessible chromatin coupled to STARR-seq (ATAC-STARR-seq), have combined STARR-seq with assays that capture accessible chromatin to extract and specifically assay the regulatory potential of accessible DNA (Buenrostro et al. 2013; Wang et al. 2018; Chaudhri et al. 2020; Glaser et al. 2021). In this way, these assays only sample a fraction of the human genome (∼2%) while assaying nearly all regulatory elements capable of driving transcription endogenously, because they are derived from open chromatin. This approach remains comprehensive of nearly all active elements and enables deeper sequencing coverage of biologically relevant regulatory elements.

Here we demonstrate an updated ATAC-STARR-seq workflow that substantially expands the capabilities of these prior methods. In addition to regulatory activity, our critical improvements enable simultaneous profiling of chromatin accessibility and TF footprinting. These additional data enable stratification of regulatory elements into TF-defined gene regulatory networks that regulate distinct cellular processes. Furthermore, the improvement in coverage we obtain allows us to identify and characterize silent regulatory regions, which are not typically identified with standard STARR-seq approaches and represent an underappreciated aspect of gene regulatory activity.

In summary, our key experimental and analytical improvements to ATAC-STARR-seq enable quantification of three complementary components of gene regulation—chromatin accessibility, TF occupancy, and regulatory activity. With this improved workflow, we capture a highly-layered multi-omic view of the human genome from a single dataset—a feature that has not been reported by regulatory assays previously. Importantly, we provide a protocol and code repository so that our new ATAC-STARR-seq workflow may be easily used and adopted by the field.

## RESULTS

### ATAC-STARR-seq Experimental Design

The ATAC-STARR-seq approach is divided into the three main parts: 1) ATAC-STARR-seq plasmid library generation, 2) reporter assay, and 3) data analysis (Figure 1A). To generate ATAC-STARR-seq plasmid libraries, nuclei are isolated from a cell type of interest and exposed to Tn5, the cut-and-paste transposase used in the ATAC-seq method (Buenrostro et al. 2013). Tn5 simultaneously cleaves DNA fragments within accessible chromatin and attaches customizable sequence adapters to their 5’ ends. ATAC-STARR-seq adapters are designed to serve as homology arms for direct Gibson cloning into the STARR-seq reporter plasmid, which enables cloning of accessible DNA fragments *en masse*. The resulting ATAC-STARR-seq plasmid library consists of millions of unique plasmids each harbouring their own unique open chromatin-derived DNA fragment.

**Figure 1.**
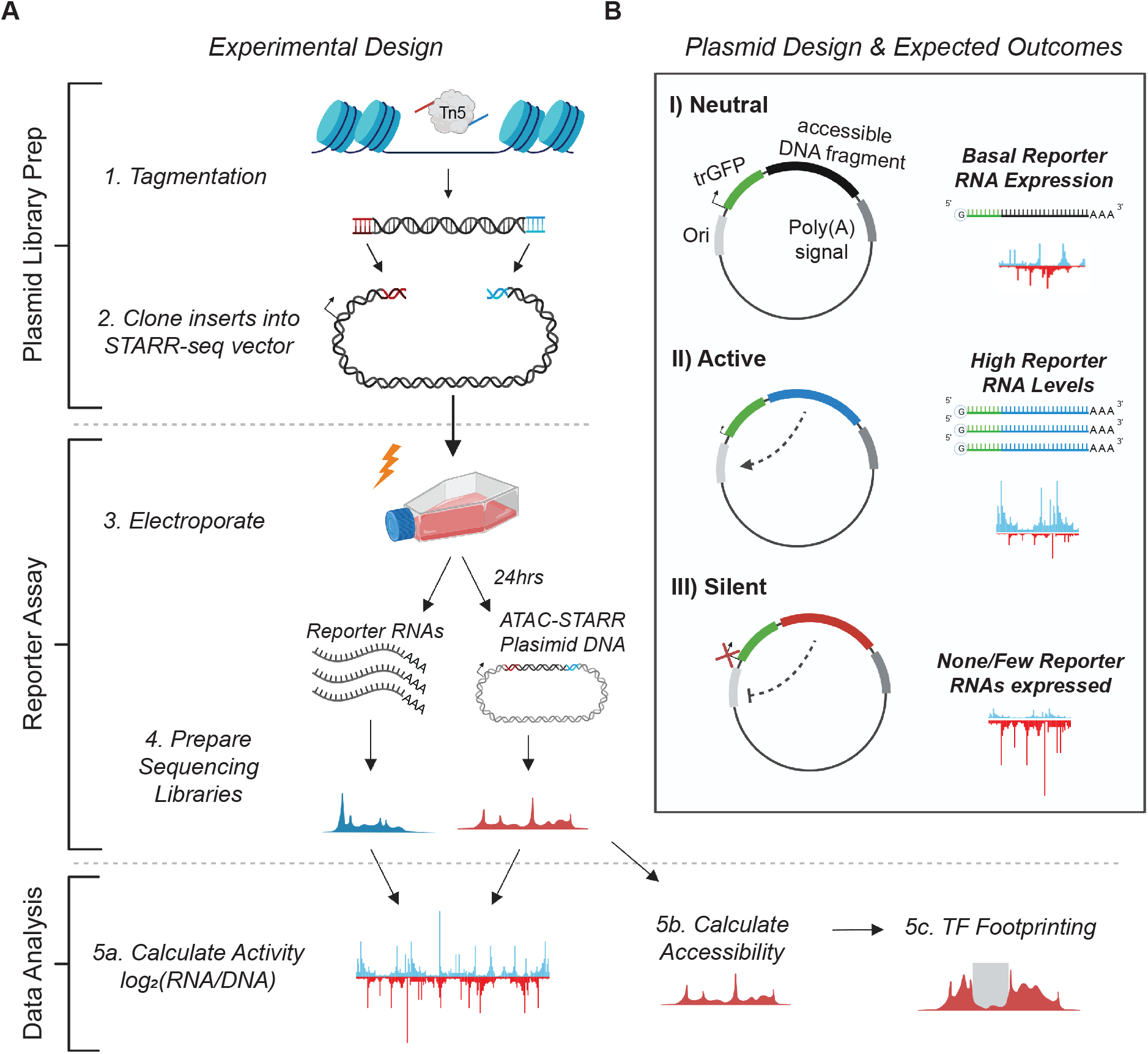
Schematic of the ATAC-STARR-seq methodology. (A) The experimental design of ATAC-STARR-seq consists of three parts: plasmid library generation, reporter assay, and data analysis. Open chromatin is isolated from cells with the cut and paste transposase Tn5. The open chromatin fragments are cloned into a reporter plasmid and the resulting clones—called an ATAC-STARR-seq plasmid library—are electroporated into cells. 24 hours later, both reporter RNAs (blue)—which are transcribed directly off the ATAC-STARR-seq plasmid—and ATAC-STARR-seq plasmid DNA (red) are harvested, and Illumina-sequencing libraries are prepared and sequenced. The resulting ATAC-STARR-seq sequence data is analysed to extract regulatory activity, chromatin accessibility, and transcription factor footprints. (B) Reporter plasmid design and the expected outcomes for neutral, active, and silent regulatory elements. Each ATAC-STARR-seq plasmid within a library contains a truncated GFP (trGFP) coding sequence, a poly-adenylation signal sequence, an origin of replication (Ori) (which moonlights as a minimal core promoter), and the unique open chromatin fragment being assayed. Since the accessible region is contained in the 3’ UTR, the abundance of itself in the transcript pool reflects its activity. In this way, neutral elements do not affect the system and reporter RNAs are expressed at a basal expression level dictated by the minimal core promoter, the Ori. Accessible chromatin fragments that are active express reporter RNAs at a higher level than the basal expression level, while silent elements repress the Ori and reporter RNAs are expressed at a lower level than basal expression.

In our updated ATAC-STARR-seq workflow, we employ the STARR-seq Ori backbone, where the origin of replication (Ori) functions as the minimal promoter (Muerdter et al. 2018). Each plasmid in the ATAC-STARR-seq plasmid library contains a truncated GFP (trGFP) coding sequence, a poly-adenylation signal sequence, the Ori, and the unique accessible DNA fragment being assayed (Figure 1B). Critically, the accessible region is cloned into the 3’ UTR, so if the accessible region is active, it interacts with the Ori to drive self-transcription. Thus, an accessible region’s level of activity is reflected by its own level of expression. Importantly, transcripts from ATAC-STARR-seq plasmids, termed “reporter RNAs”, are expressed at basal levels from the activity of the Ori itself. This allows detection of silencing activity—the inhibition of the basal expression—in this assay.

Following its creation, the ATAC-STARR-seq plasmid library is transfected via electroporation into a given cell line. From the same flask of cells, both reporter RNAs and plasmid DNA are harvested 24 hours later, then prepared as Illumina sequencing libraries and sequenced. Activity is calculated as the log2 ratio between normalized read counts from the reporter RNA and plasmid DNA datasets. Notably, the re-isolation of plasmid DNA recovers only the ATAC-STARR-seq plasmids that were successfully transfected, thus providing a more accurate representation of the “input” sample than sequencing without transfection.

### ATAC-STARR-seq maintains library complexity and nucleosome profiles of Tn5 selected DNA fragments

Following the experimental design outlined above, we tagmented GM12878 cells and generated an ATAC-STARR-seq plasmid library that yielded about 50 million unique accessible DNA fragments (Supplemental Text, Supplementary Figure 1A). We then transfected the library into GM12878 cells and harvested both reporter RNAs and plasmid DNA from the same flask of cells 24 hours later. We chose 24 hours post-transfection to avoid significant effects from the plasmid-induced interferon gene response, and to ensure the data reflects steady-state regulatory properties of GM12878 accessible regions (Supplemental Text, Supplementary Figure 1B) (Muerdter et al. 2018). Using the captured reporter RNAs and plasmid DNA, we prepared illumina sequencing libraries and submitted for sequencing. In total, we performed three replicates.

The size distribution of the accessible DNA fragments remained remarkably consistent throughout the ATAC-STARR-seq procedure and displayed the characteristic nucleosome banding and DNA pitch typified by ATAC-seq fragment libraries (Supplementary Figure 2A-B). Analysis of library complexity between replicates revealed an average maximum complexity of 90 million unique fragments for input DNA, and 10 million unique fragments for reporter RNAs (Supplementary Figure 2C). The difference between RNA and DNA complexities is likely due to higher duplication rates in the RNA samples (Supplementary Table 1) driven by both the expression of multiple transcripts per plasmid and more PCR cycles required for the RNA samples. In addition, for both RNA and DNA samples, replicates displayed high pearson (r^2^: 0.96-0.99) and spearman (ρ: 0.77-0.93) correlation coefficients indicating strong agreement among the three replicates assayed (Supplementary Figure 3). Altogether the ATAC-STARR-seq sequence libraries demonstrated the necessary quality and complexity for downstream analysis.

### ATAC-STARR-seq faithfully captures chromatin accessibility with high signal-to-noise

The use of Tn5 on native chromatin to selectively clone chromatin accessible DNA fragments provides the opportunity to quantify not only reporter activity, but also chromatin accessibility simultaneously from the same plasmid library. However, ATAC-STARR-seq involves many additional steps than standard ATAC-seq, such as cloning and transfection, which might distort the chromatin accessible landscape. To validate the robustness of ATAC-STARR-seq to measure accessibility, we called accessibility peaks using data from the three ATAC-STARR-seq reisolated plasmid DNA replicates and compared against the original GM12878 ATAC-seq dataset from Buenrostro *et al*. 2013. Raw sequences obtained for both datasets were processed through identical workflows (see Methods). After collapsing read duplicates, peaks were called at a false-discovery rate (FDR) of 0.01%, which produced a peak count similar to the number previously reported for the Buenrostro *et al*. 2013 data (∼73,000) (peak counts for a variety of FDR cut-offs are supplied in Supplementary Table 2). The ATAC-STARR-seq and Buenrostro peak sets represent 2.22% and 1.74% of the genome, respectively, which agrees with previous reports (Figure 2A) (Thurman et al. 2012; Klemm et al. 2019). Overall, 61% of ATAC-STARR-seq peaks are reproduced in the Buenrostro dataset, while 86% of Buenrostro peaks overlap the ATAC-STARR-seq dataset (Figure 2B; Jaccard index = 0.48), indicating strong agreement between these data despite substantial differences in ATAC-STARR DNA sample preparation. Based on this, we conclude that ATAC-STARR-seq can accurately retain chromatin accessible peaks in the human genome.

**Figure 2.**
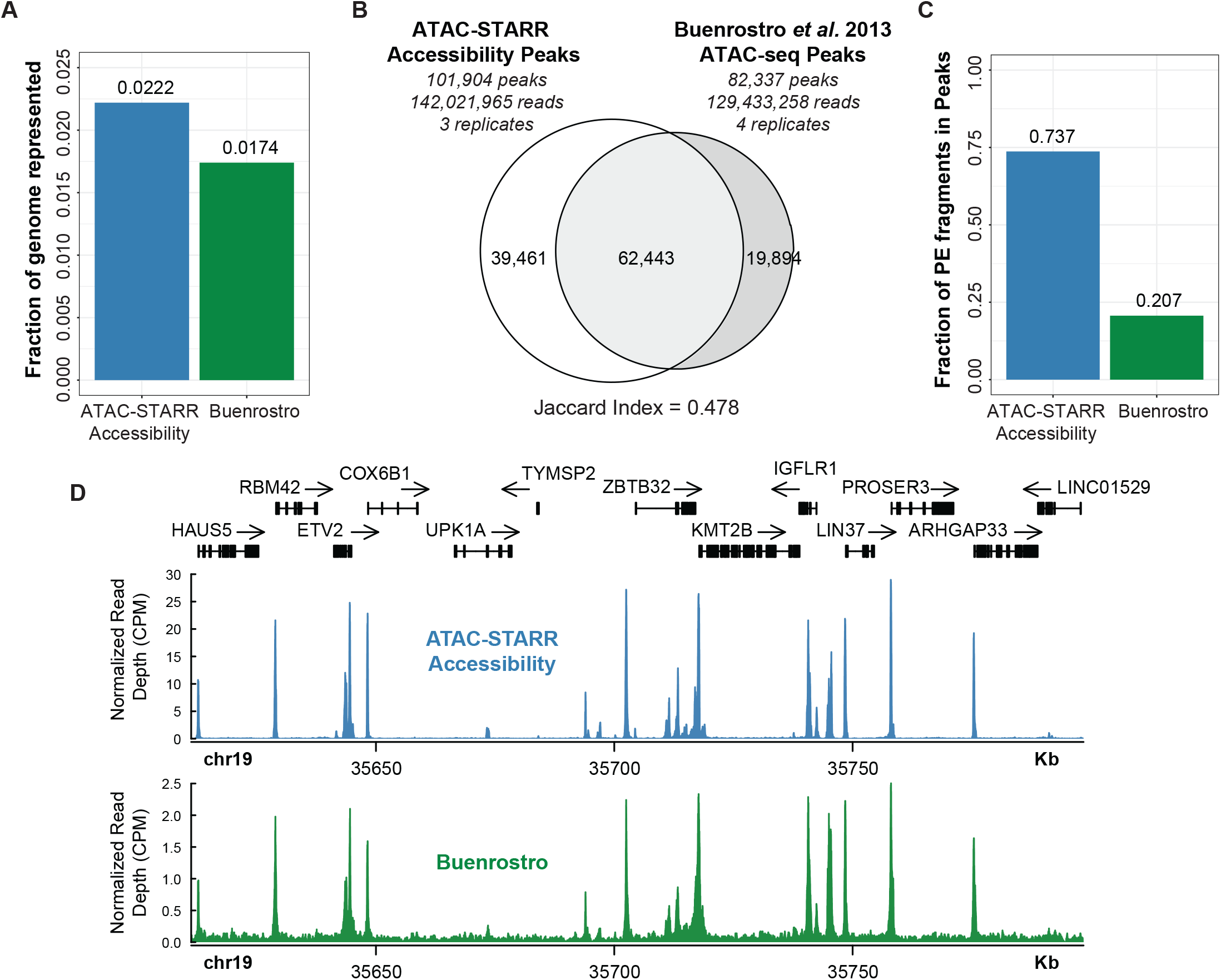
ATAC-STARR-seq accurately quantifies chromatin accessibility. ATAC-seq data from Buenrostro et al. 2013 is compared with ATAC-STARR-seq plasmid DNA data. (A) Fraction of the human genome represented by each peakset. (B) Venn diagram of peak overlap between the two datasets and the associated Jaccard Index. (C) Fraction of paired-end (PE) fragments in peaks—FRiP scores—for both samples. (D) Signal tracks comparing counts per million (CPM) normalized read count at a representative locus.

While peak sets are highly similar, we observe a notable difference in background signal. The fraction of reads in peaks score (FRiP), an ENCODE ATAC-seq standard measure of noise, is considerably higher for ATAC-STARR-seq (0.74) than observed for Buenrostro (0.21) or the ENCODE accepted standard (>0.2) (Figure 2C). This dramatic difference in FRiP indicates that ATAC-STARR-seq provides significantly less background noise or spurious reads than a typical ATAC-seq library. One possible explanation is that each plasmid library is much more complex than a standard ATAC-seq experiment, consisting of a pool of eight individual Tn5 tagmentation reactions per replicate compared to one. Another is the use of the Omni-tagmentation approach, which is known to reduce background and improve FRiP scores (Corces et al. 2017). This difference is also seen when comparing signal at a representative locus (Figure 2D). With counts per million normalized samples, we observe dramatically different peak intensities and see very little background signal in the ATAC-STARR-seq sample. Thus, ATAC-STARR-seq captures chromatin accessibility with high signal-to-noise.

### A sliding windows approach increases activity region calling sensitivity

ATAC-STARR-seq tests regulatory activity in DNA enriched for accessible chromatin. Unlike whole genome STARR-seq or other MPRAs, where the genomic DNA fragment distribution is relatively constant, read coverage varies dramatically from peak-to-peak in ATAC-STARR-seq. In this way, ATAC-STARR-seq requires an analysis strategy that calls active and silent regulatory regions within accessibility peaks. To address this “peaks-within-peaks” problem, we developed an analytical approach using DESeq2 to normalize reporter RNA read counts to reisolated plasmid DNA read counts. DESeq2 additionally performs an independent filtering step which removes low count data confounders that can dramatically influence ratios and result in false positive peak calls (Love et al. 2014).

We tested two different approaches for regulatory activity analysis. The two approaches differ in how genomic regions are defined prior to differential analysis with DESeq2. Our “sliding window” method, defines regions by slicing accessible peaks into 50bp sliding bins with a 10bp step size (Figure 3A). Alternatively, the “fragment group” method, which is the approach used in Wang *et al*. 2018, synthesizes regions by grouping paired-end sequencing fragments by 75% or greater overlap (Supplementary Figure 4A). Using a different set of genomic regions, both methods assign and count overlapping RNA and DNA reads to each genomic region and, using DESeq2, identify regions where the RNA count is statistically different from the DNA count at a Benjamini-Hochberg (BH) adjusted p-value < 0.1. The “sliding window” method yielded ∼30,000 distinct active regions, while the “fragment groups” method yielded ∼20,000 distinct active regions (Supplementary Figure 4B). In addition, nearly all active regions defined using the fragment group method (95%) are also captured in the sliding window method regions (Supplementary Figure 4C). Given this overlap and a 50% greater recovery with the sliding windows approach, we used the sliding windows method to call active ATAC-STARR-seq regulatory regions.

**Figure 3.**
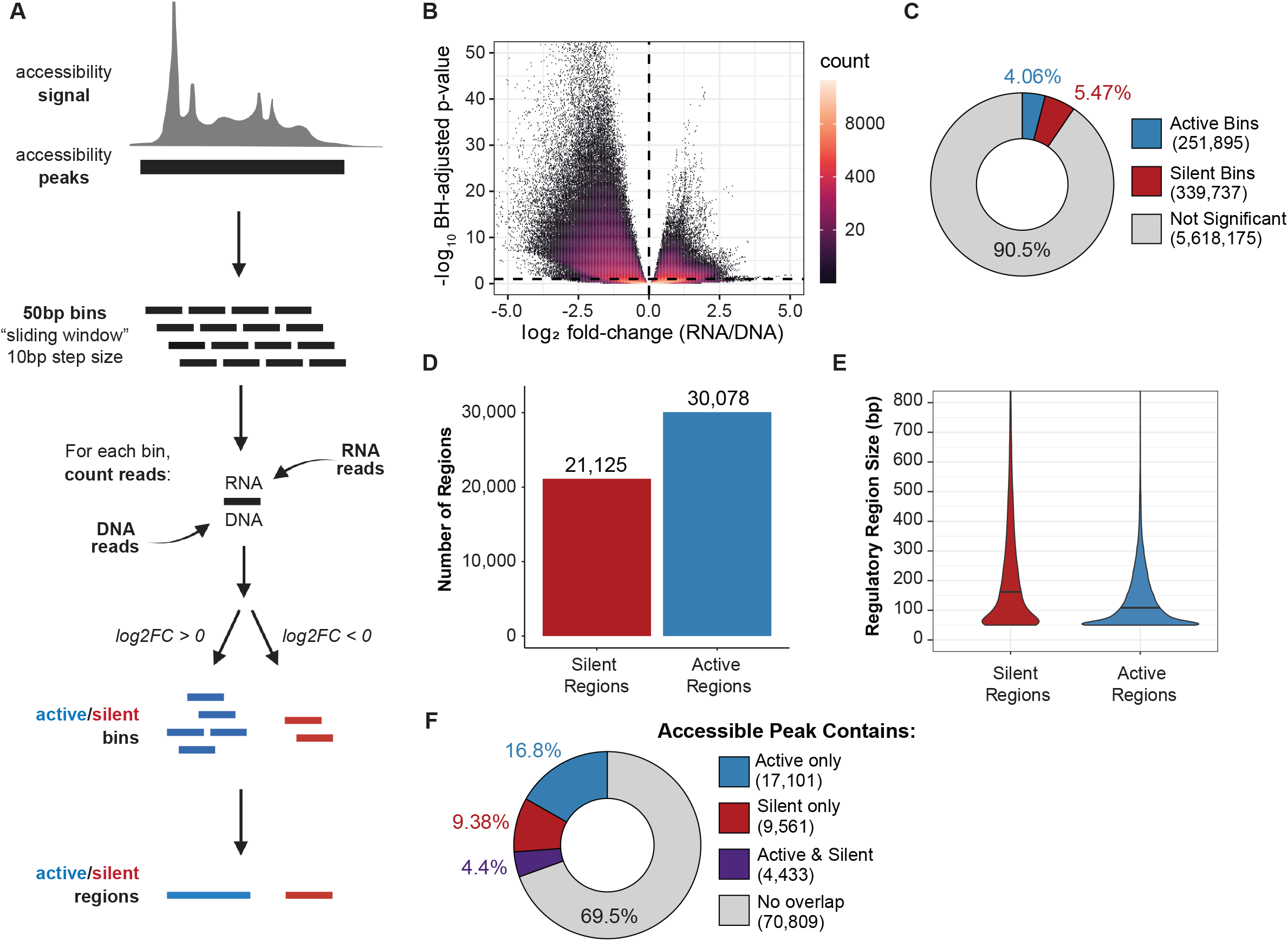
ATAC-STARR-seq quantifies regulatory activity within accessible chromatin. (A) Schematic of the sliding window peak calling method. Accessibility peaks are chopped into 50bp bins at a 10bp step size with BEDtools’ makewindows function (opons -w 50, -s 10). For each window, RNA and DNA reads are counted using Subread’s featureCounts function. Differential analysis comparing RNA and DNA read count is performed with DESeq2. Significant bins are called at an Benjamini-Hochberg (BH) adjusted p-value < 0.1 and parsed into active or silent depending on log2 fold-change (FC) value (+/- zero). Finally, bins are collapsed into regions using BEDtools’ merge function. Log2FC scores are averaged across merged bins. (B) Volcano plot of log2FC scores against -log10-transformed BH adjusted p-value from DESeq2 for all bins analyzed. (C) The proportion of bins called as active or silent. (D) The number of regions defined as either active or silent. (E) A violin plot of active and silent regulatory region size; solid lines represent the mean in each case. (F) The proportion of accessible peaks that overlap an active or silent region, or both.

Because significance is the primary threshold in our region calling strategy, we examined the influence of replicate count on the number of active regions called (Supplementary Text, Supplementary Figure 5). We found that, as expected, more replicates result in more active regions. However, we caution that these additional regions may represent a disproportionate number of false positives and may affect the outcomes of certain accuracy-sensitive applications like computational modelling. In this way, we believe three replicates are sufficient for most purposes. We also investigated the effect of removing read duplicates and found that it substantially hindered sensitivity (Supplementary Text, Supplementary Figure 6).

### ATAC-STARR-seq quantifies regulatory activity of open chromatin

In the sliding window approach, bins are classified as active or silent depending on whether RNA is enriched or depleted, respectively, and then like-bins are merged to collapse overlaps (Figure 3A). Using this approach, we identified ∼590,000 bins where RNA and DNA counts were significantly different (Figure 3B). More specifically, this analysis identified 251,895 (4.1%) active bins and 339,737 (5.5%) silent bins from the ∼5.6 million total bins measured (Figure 3C). Overlapping bins were merged into 30,078 active and 21,125 silent regulatory regions (Figure 3D). Notably, we see more silent than active bins called but, because silent regions are generally larger (Figure 3E), merging overlapping bins results in fewer silent regions than active. Collectively, the active and silent bins represent ∼9.5% of all bins measured, indicating that the majority of accessible DNA is transcriptionally neutral. Moreover, most accessible peaks do not have an active or silent region contained within them (69.5%), suggesting that most accessible regions are neutral regulatory regions according to our assay (Figure 3F). This suggests that the majority of accessible DNA has no regulatory potential in this context or, alternatively, that ATAC-STARR-seq is not sensitive enough to measure weakly active or weakly silent regions. A recent study in mouse embryonic stem cells made the same observation using an orthogonal approach, suggesting this phenomenon is preserved in other mammalian species (Glaser et al. 2021). Interestingly, a small percentage of accessible peaks (4.4%) contain both active and silent regions, demonstrating that there can be competing regulatory regions within the same accessible peak.

### Active and silent ATAC-STARR-seq regions represent both proximal and distal *cis*-regulatory elements and lie within functional chromatin states

To gain insight into the regulatory features of active regions, we annotated both active and silent regions according to genomic location. Active regions are found in both promoter proximal and distal areas of the genome, with a majority occurring in intronic and intergenic sites (∼55%), whereas silent regions coincide primarily with promoters (∼75%) (Figure 4A). Functional classification of active and silent regions by the 18-state ChromHMM model (Roadmap Epigenomics et al. 2015) revealed that active regions consist of TSS active, TSS flanking upstream, and Enhancer Active 1 chromatin states and are devoid of repressive states like Repressed Polycomb Weak and Quiescent (Figure 4B). By contrast, silent regions are slightly enriched for bivalent chromatin states (TSSBiv, EnhBiv), consistent with the observation that they are accessible but not active. Interestingly, most silent regions also coincide with TSS Active and TSSFlnk ChromHMM states, which corroborates their promoter proximal locations; however, their designation as “active” by chromHMM is somewhat puzzling considering these DNA fragments do not drive transcription in our assay. One explanation is that silent regulatory activity, as measured by episomal-based reporter assays, does not fully copy regulatory activity as predicted by ChromHMM. Alternatively, active promoters may confound the reporter assay by initiating transcription from the 3’UTR of the plasmid causing conflicts with active transcription from the Ori.

**Figure 4.**
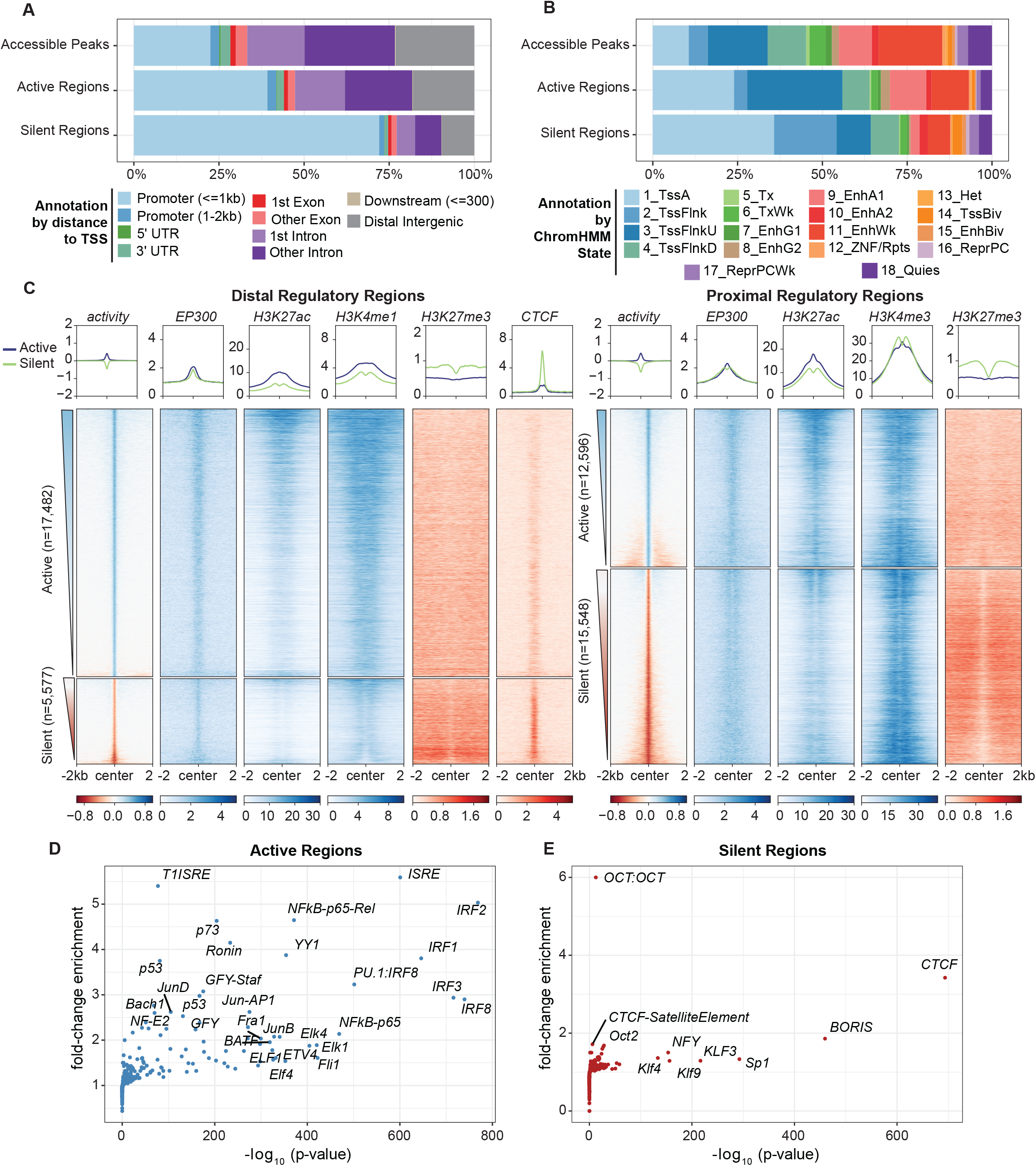
Regulatory regions defined by ATAC-STARR exhibit annotations, histone modifications, and TFs characteristic of their function. (A) Annotation of regulatory regions relative to the transcriptional start site (TSS). The promoter is defined as 2kb upstream and 1 kb downstream of the TSS. (B) Annotation of regulatory regions by the ChromHMM 18-state model for GM12878 cells. (C) Heatmaps of GM12878 ENCODE ChIP-seq signal and regulatory activity for proximal and distal ATAC-STARR-defined regulatory regions. Proximal regions were classified as within 2kb upstream and 1 kb downstream of a TSS; all other regions were annotated as distal. Active and silent regions were ranked by mean activity signal for both proximal and distal regions. (D-E) Transcription factor motif enrichment analysis as quantified by HOMER. Fold-change values are relative to the default background calculated by HOMER.

To further investigate if silent regions are a result of 3’UTR transcription initiation, we considered if an orientation bias existed in reporter RNAs levels. If 3’UTR transcription conflicts exist, we would expect many fewer reporter RNAs when this transcription results in head-on conflicts rather than occurring in the same direction as the Ori. We therefore subset reads based on whether they arose from an insert cloned in a 3’ to 5’ direction or in a 5’ to 3’ direction (Supplementary Figure 7A). We then assigned read counts to all bins analyzed (Supplementary Figure 7B-C), the bins called active (Supplementary Figure 7D-E), or the bins called silent (Supplementary Figure 7F-G). Because this is expected to be a promoter-specific effect, we also split bins into proximal and distal based on location to the nearest transcription start site. In all cases, more than 95% of the bins do not display an orientation bias, which we defined as a normalized read count difference greater than five between orientations (Supplemental Methods, Supplementary Figure 7H). Moreover, we observe high pearson and spearman correlation coefficients between orientations for all conditions (r^2^: 0.80-0.91 and ρ: 0.73-0.90) and the minimal contribution of orientation bias to silent regions is in agreement with a previous report (Klein et al. 2020). For the <5% of regions that do display orientation bias, proximal bins are more affected than distal bins, as expected. Altogether, ATAC-STARR-seq does not display a significant orientation bias and most of the 21,000 silent regions we observe result from legitimate silencing activity or another source.

### Active and silent ATAC-STARR-seq regions are distinct functional classes and are enriched for specific histone modifications and TF motifs

To further investigate the chromatin landscape of our active and silent regions, we plotted ENCODE GM12878 ChIP-seq signal (Consortium et al. 2020) within active and silent regions for EP300, CTCF, and histone modifications associated with active and repressed states (Figure 4C). As expected, active regions contain EP300 at their center with histone 3 lysine 27 acetylation (H3K27ac) more broadly distributed across the center; histone 3 lysine 4 mono-methylation (H3K4me1) is also present at distal regions, while histone 3 lysine 4 tri-methylation (H3K4me3) is at proximal regions. In addition, histone 3 lysine 27 tri-methylation (H3K27me3)—a bivalent repressive mark—is largely absent from active regions. Proximal silent regions, on the other hand, are enriched for H3K27me3 and H3K4me3. This suggests many of the proximal silent regions may be accessible bivalent regulatory elements in lymphoblastoid cells. Further, distal silent regions are strongly enriched for CTCF, suggesting many of these sites may be CTCF-bound insulators. To support their designation as silent calls, we also compared histone modification signal at accessible peaks that contain either a silent region, an active region, both a silent and active region, or neither, which we define as neutral accessible peaks (Supplementary Figure 8). Consistent with the observations above, silent accessible peaks contain more H3K27me3 signal and are devoid of H3K27ac signal relative to the other accessible peak types.

We also performed motif enrichment analysis and observed widely different sets of TF motifs enriched in either the active or silent regions (Figure 4D-E). The most enriched TFs in active regions were the IRF family, the ETS family, NFkB, and general promoter TFs such as Ronin/GFY and YY1. Silent regions are enriched for CTCF and its counterpart BORIS, which are well characterized insulator proteins and are typically transcriptionally inert. Beyond insulators, we found enrichment for the SP/KLF family several of which are known to be transcriptional repressors (Cao et al. 2010). These data are consistent with our current understanding of immune gene regulation and regulatory element function, which together corroborates the quantification of regulatory activity with ATAC-STARR-seq.

### ATAC-STARR-seq retains the ability to map *in vivo* TF binding

An inherent advantage of an ATAC-seq based approach is the ability to perform TF footprinting. Computational footprinting methods identify Tn5 cleavage events or “cut sites” from ATAC-seq data and, when combined with motif analysis, can identify TF binding sites with high accuracy (Bentsen et al. 2020; Yan et al. 2020). Since ATAC-STARR-seq produces similar chromatin accessibility peak sets as standard ATAC-seq, we explored whether TF footprints were also preserved. We generated Tn5-bias corrected cut site signal files for both Buenrostro and ATAC-STARR-seq accessibility datasets and plotted cut site signal at all accessible CTCF motifs (Figure 5A) (Bentsen et al. 2020). As a control, we also plotted GM12878 CTCF ChIP-seq signal from ENCODE and ranked region order by highest mean ChIP-seq signal. We observed consistent depletion of Tn5 cut-sites for both Buenrostro and ATAC-STARR-seq accessibility at CTCF sites. Moreover, we only observe footprints at motifs with CTCF ChIP-signal, demonstrating the utility of TF footprinting to determine motifs that are bound or unbound by TFs. Given the importance of TFs in driving enhancer function, this distinction is vital when dissecting transcriptional regulation in human cells.

**Figure 5.**
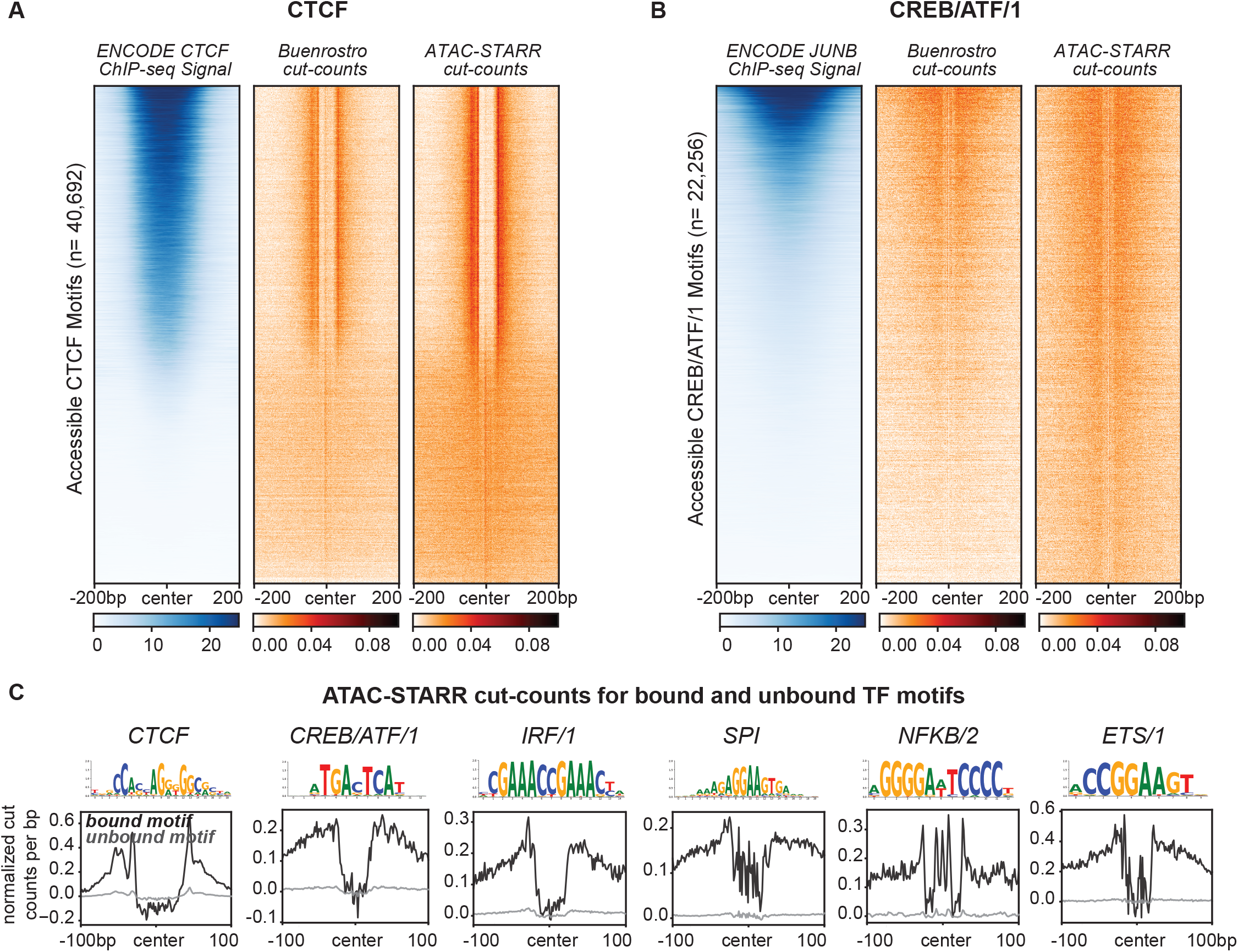
ATAC-STARR-seq identifies transcription factor footprints. (A) Comparison of ENCODE CTCF ChIP-seq signal to Buenrostro and ATAC-STARR-seq cut count signal for all accessible CTCF motifs. (B) Comparison of ENCODE JUNB ChIP-seq signal to Buenrostro and ATAC-STARR-seq cut count signal for all accessible motifs with the CREB/ATF/1 motif archetype. For both, regions were ranked by largest mean ChIP-seq signal. (C) Aggregate plots representing mean signal for the TOBIAS-defined bound and unbound motif archetypes: CTCF, CREB/ATF/1, IRF/1, SPI, NFKB/2, ETS/1.

TF motif enrichment pointed to multiple CREB/ATF family members, including JUNB which is a critical immune cell regulator (Figure 4D). So, we asked whether JUNB footprints are also present in our data. Unlike CTCF, JUNB shares its motif with many other transcription factors, such as ATF1; therefore, footprinting cannot distinguish JUNB and ATF1 binding sites. For this reason, we refer to TFs using their ENCODE-defined “archetypes”, which reflects the group of TFs that share the same motif (Vierstra et al. 2020). For each archetype, we performed footprinting against one of the TFs within an archetype to infer motifs bound by members of the group, such as JUNB for the CREB/ATF/1 archetype. To assess the extent to which JUNB footprints can be explained by JUNB binding, we plotted GM12878 JUNB ChIP-seq signal from ENCODE within both Buenrostro and ATAC-STARR-seq cut sites (Figure 5B). Indeed, JUNB ChIP-seq signal explains some but not all the CREB/ATF/1 footprints present. Importantly, we observe similar cut-site signal to Buenrostro, further indicating that ATAC-STARR-seq can detect *in vivo* binding of transcription factors despite the additional cloning and transfection steps involved in producing ATAC-STARR-seq DNA libraries.

We footprinted against several more TF archetypes to identify bound or unbound TF motifs (Figure 5C). For all TFs, bound motifs display dramatically larger footprint depth than unbound motifs. Together, this indicates that ATAC-STARR-seq, when combined with footprinting, can identify regions of the genome where TFs are bound. This additional level of information can be leveraged in conjunction with accessibility and activity to understand the context of TF binding while circumventing the need to perform individual chromatin immunoprecipitations.

### Collective profiling of accessibility, *in vivo* TF binding, and activity with ATAC-STARR-seq reveals distinct networks of gene regulation

Interrogating chromatin accessibility, TF binding, and regulatory activity together can be used to interpret locus-specific gene regulatory mechanisms. For example, active regulatory elements surrounding the B cell-specific expressed gene *ZBTB32* overlap JUNB footprints suggesting these regions are regulated by JUNB binding (Figure 6A). We also observe a CTCF footprint overlapping a silent region at the *ETV2* locus, a gene lowly expressed in B cells, according to the Human Protein Atlas (Uhlen et al. 2015; Uhlen et al. 2019). Together this indicates that active and silent regions can, in part, be explained by the occupancy of these TFs.

**Figure 6.**
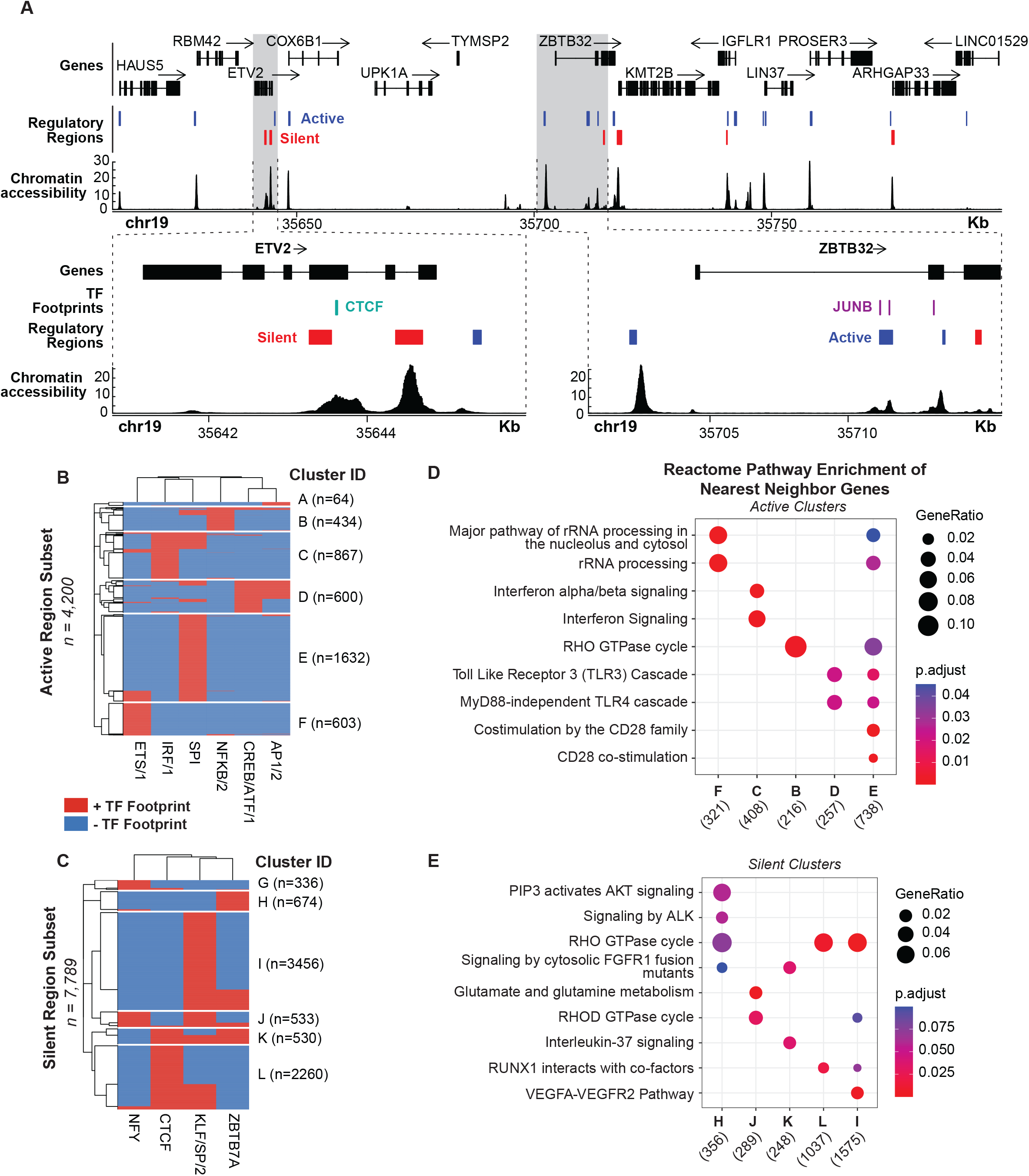
TF footprints stratify ATAC-STARR-defined regulatory regions into gene regulatory networks. (A) ATAC-STARR-defined chromatin accessibility, TF footprints, and regulatory regions at chr19:35,611,232-35,798,446 (hg38). Signal tracks represent counts per million normalized read depth of chromatin accessibility. Zooms into ETV2 and ZBTB32 show that some regulatory regions are occupied by a CTCF or JUNB footprint. (B-C) Heatmaps of clustered (B) active and (C) silent regions based on presence or absence of footprints for select TF motif archetypes. (D-E) Reactome pathway enrichment analysis for nearest-neighbor gene sets for each of the clusters. Genes counts for each cluster are displayed below their ID.

To demonstrate the power of integrating TF footprints and regulatory regions on a global scale, we clustered active and silent regions based on the presence or absence of several TF footprints (Figure 6B-C). Footprints were selected based on top hits from the previous motif enrichment analysis (Figure 4D-E). Regulatory activity may be driven by one or multiple TF binding events that defines the cluster and is representative of a gene regulatory network in the genome. Indeed, we find that the putative target genes regulated by each unique group are enriched for distinct gene regulatory pathways and are often related to the TFs in the cluster (Figure 6D-E). For example, cluster C is primarily defined by the presence of IRF/1 and is enriched for interferon alpha/beta signalling. It is interesting that active clusters tend to be more associated with B cell function than silent clusters, which are more associated with non-B cell related pathways.

Altogether, these distinct gene regulatory networks provide an additional layer of insight into the mechanisms that control gene expression and showcase how integration of the multi-omic information provided by ATAC-STARR-seq can narrow our focus of gene targets for active and silent regions. We envision such an analysis could be used to interpret the functional consequences of a dysregulated transcription factor or disease-associated genetic variants. Importantly, we provide this level of detail from a single dataset, which further demonstrates the strong potential of our workflow to reveal distinct functional layers of human gene regulation. The resolution we achieve here not be possible without all three levels of regulatory information provided by ATAC-STARR-seq.

## DISCUSSION

Genome-wide approaches that integrate measurements of multiple layers of gene regulation are needed to better understand enhancer function. By combining ATAC-seq with STARR-seq, ATAC-STARR-seq assays regulatory activity only within the context of accessible chromatin. This allows deeper coverage of regulatory elements by narrowing scope but remaining inclusive of nearly all active regulatory elements. In this report, we substantially expand the capabilities of ATAC-STARR-seq and present an improved workflow which uniquely permits simultaneous profiling of accessibility, TF occupancy, and regulatory activity from a single DNA fragment source. Specifically, we implement key experimental and analytical improvements and present data rationalizing the decisions we make. Experimentally, we adapt a modified tagmentation protocol to remove mitochondrial DNA from the DNA fragment pool. We also utilize the Ori as the minimal promoter on our STARR-seq backbone which improves reporter RNA expression, recovery, and dynamic range over the super core promoter (SCP1) backbone (Muerdter et al. 2018; Klein et al. 2020). Furthermore, we reisolate the transfected plasmid DNA to capture only the DNA that is available to cells, which is a more accurate measure of the input than sequencing prior to transfection. Moreover, reisolating plasmid DNA drives a greater degree of variance between samples and better reflects a replicate than sequencing the same DNA sample for each replicate. Critically, we also developed and tested a simple and sensitive region calling strategy that improves detection of regulatory regions including silencers. We also quantify chromatin accessibility and identify TF footprints, which is surprising given the added processing steps in ATAC-STARR-seq including cloning, transfection, and recapture of DNA libraries that would dull or degrade footprint signal. This enabled us to stratify the active and silent regulatory regions into distinct gene regulatory networks defined by the presence of one or multiple TF footprints. Such an analysis typically requires multiple sequencing assays, but we do this using a single dataset.

With this improved workflow, we identified 30,078 active regions and 21,125 silent regions in lymphoblastoid cells. Most active regions were distal to transcription start sites, enriched for functional active ChromHMM states, and were enriched for known B cell regulating-TF motifs such as IRF8 and NFkB. By contrast, the silencers are proximal to transcription start sites and enriched for CTCF and the SP/KLF TF family. Silent regions are also enriched for the bivalent marks H3K27me3 and H3K4me3 and may represent regulatory regions that are poised, particularly at promoters. Because our plasmid design places regulatory regions within the 3’UTR of the truncated reporter gene, it is possible that the lack of observed reporter RNAs at silent regions are a result of head-on transcriptional conflicts that arises from antisense transcription initiation from the 3’UTR. However, we show this minimally occurs in our system and the silent regions reflect true silencing activity or another source that has yet to be identified. While further studies may be needed to validate these silent regions, this work confirms that the silencers are a distinct class of regulatory element with distinct properties compared to active and neutral regions and warrant further investigation. Even with an increasing number of studies targeted at identifying silencers in the human genome, silencing regulatory regions remain an under-studied aspect of gene regulation and our approach provides a new strategy to investigate these elements on a global scale (Doni Jayavelu et al. 2020; Pang and Snyder 2020; Kim et al. 2021).

ATAC-STARR-seq data has several distinct attributes that require a tailored analysis strategy. Current MPRA bioinformatic tools and pipelines are not tractable for these data, because in ATAC-STARR-seq the input itself is enriched for accessible chromatin and the read pileup varies dramatically within these loci. In this way, the analysis of our data required us to essentially call “peaks within peaks”. For this reason, it was critical to 1) normalize RNA to DNA and 2) avoid regions of low count data, which is why we adapted approaches using DESeq2. We also showed that including PCR duplicates was preferred over collapsing duplicates. In the future it would be beneficial to introduce a unique molecular identifier to the system—such as the strategy employed by UMI-STARR-seq (Neumayr et al. 2019)—to collapse only the duplicates arising from PCR. While we show comparisons of analysis strategies here, we believe that more information could be extracted from this and future ATAC-STARR-seq datasets with improved analysis strategies. In recent years we have seen the development of tailormade peak callers for whole genome STARR-seq, such as CRADLE (Kim et al. 2021) and STARRPeaker (Lee et al. 2020); a similarly tailored ATAC-STARR-seq peak caller could further improve the capabilities of the method.

While this study was limited to one condition, there are many potential applications of ATAC-STARR-seq. With the ability to subset enhancers by TF binding, ATAC-STARR-seq could be leveraged to investigate enhancer grammar by pairing measurable regulatory activity with multiple TF footprints that drive enhancer function. This approach also has the potential to identify dysfunctional gene regulatory networks in diseases like cancer where neoplastic transformation can be driven by the dysfunction of a specific TF. Additionally, an ATAC-STARR-seq plasmid library may be generated from one cell-type and tested in another. This flexibility could be used as a tool to determine context dependent activity or investigate enhancer and TF usage patterns during a differentiation time course.

In this study, we demonstrated that our improved ATAC-STARR-seq workflow is a powerful approach enabling joint quantification of chromatin accessibility, transcription factor occupancy, and regulatory activity. We further demonstrate how this single assay can characterize the human genome at many functional levels from chromatin accessibility to distinct gene regulatory networks. This method provides a state-of-the-art approach to deeply investigate transcriptional regulation of the human genome. We provide a detailed protocol and a well-documented code repository so that ATAC-STARR-seq may be easily used and adapted by the field.

## MATERIAL AND METHODS

### Cell Culture

GM12878 cells were obtained from Coriell and cultured with RPMI 1640 Media containing 15% fetal bovine serum, 2mM GlutaMAX, 100 units/mL penicillin and 100 μg/mL streptomycin. Cells were cultured at 37°C, 80% relative humidity, and 5% CO2. Cell density was maintained between 0.2×10^6^ and 1×10^6^ cells/mL with a 50% media change every 2-4 days. All cell lines were regularly screened for mycoplasma contamination using the MycoAlert kit (Lonza).

### Plasmids

The hSTARR-seq_ORI plasmid vector was a gift from Alexander Stark (Addgene plasmid #99296) and the pcDNA3-EGFP was a gift from Doug Golenbock (Addgene plasmid #13031). The bacterial stabs from Addgene were spread onto an LB agar plate containing 100μg/mL ampicillin and incubated at 37°C overnight. For each, a single colony was picked and grown in 50mL LB containing 100μg/mL ampicillin overnight at 37°C while shaking at 225rpm. Plasmid DNA was extracted using the ZymoPURE II Plasmid Midiprep kit (Zymo Research, #D4200).

The linear vector used in the ATAC-STARR-seq gibson cloning step was generated by a single 50μL PCR reaction using NEBNext^®^ Ultra™ II Q5^®^ Master Mix (NEB, #M0544S). While not necessary for this study, primers were designed to add the i5 barcode to the linearized vector; this allows for different ATAC-STARR-seq plasmid libraries to be pooled and tracked. Following this approach, a universal forward primer (Fwd_universal_STARR) and a reverse primer (Rev_N504_STARR) designed to add the N504 barcode were used (primer sequences are provided in Supplementary Table 3). PCR Products were purified with the Zymo Research DNA Clean & Concentrator-5 kit. DNA yield was determined by nanodrop, and purity was analysed by gel electrophoresis; the linearized vector was the only product observed on the gel.

### Tagmentation

A total of eight tagmentation reactions were performed on 50,000 GM12878 cells each. We followed a slightly modified version of the omni-ATAC approach used in Corces *et al*. 2017(Corces et al. 2017). Specifically, twice as much Tn5 than described in the protocol was used. Standard Tn5 transposase was prepared in-house following the method described in Picelli *et al*. 2014 (Picelli et al. 2014). Standard Tn5 transposome was assembled as described in Barnett *et al*. 2020 (Barnett et al. 2020) with the following oligos: Tn5_1, Tn5_2_ME_comp, and TN5MEREV. Tagmented products were pooled together and purified with the Zymo Research DNA Clean & Concentrator-5 kit (#D4013). The entire elution was split and amplified via five-10μL PCR reactions. We used NEBNext^®^ High-Fidelity 2X PCR Master Mix (#M0541S)—which is importantly not a hot-start formulation—to first extend tagments before the initial denaturation step of PCR via the following cycling parameters: 72°C 5 min, 98°C 30s; 4 cycles of 98°C 10s, 62°C 30s, 72°C 60s; final extension 72°C 2 min; hold at 10°C. Forward and reverse primer sequences, Fwd_atac-starr_tag and Rev_atac-starr_tag, are provided in Supplementary Table 3. Amplified products were purified with the Zymo Research DNA Clean & Concentrator-5 kit and then analyzed for concentration and size distribution with a HSD5000 screentape (Agilent, #5067) on an Agilent 4150 TapeStation system. After amplification, we selected PCR products less than 500bp using SPRISelect beads (Beckman-Coulter, #B23317) at a 0.6X volume ratio of beads:sample. Selection was verified using a HSD5000 screentape.

### Massively Parallel Cloning

Four 10μL gibson cloning reactions were performed with NEBuilder^®^ HiFi DNA Assembly Master Mix at a vector:insert molar ratio of 1:2. As a negative control, we performed one cloning reaction substituting tagments with nuclease-free water. Gibson products were pooled and purified via ethanol precipitation as previously described in Sambrook & Russell (Sambrook and Russell 2006); we used glycoblue (150μg/mL) as a co-precipitant. Purified gibson products were electroporated into MegaX DH10B T1R Electrocomp™ Cells (Invitrogen, # C640003) using a Bio-Rad Gene Pulser. Three electroporations for the ATAC-STARR-seq sample (and 1 for the control) were performed with the following parameters: exponential decay pulse type, 2kV, 200Ω, 25μF, and 0.1cm gap distance. Pre-warmed SOC media (1mL) was added immediately following electroporation; the three reactions were pooled and incubated at 37°C for 1 hour. We confirmed cloning success by plating a dilution series—using a small aliquot from the ATAC-STARR-seq and negative control samples—onto pre-warmed LB agar plates containing 100μg/mL ampicillin and visualizing colonies 24 hours later. The remaining ATAC-STARR-seq transformation was added directly to a 1L LB liquid culture with 100μg/mL ampicillin and grown at 37°C while shaking at 225rpm overnight. The next day, plasmid DNA was harvested from the 1L culture using the ZymoPURE II Plasmid Gigaprep (Zymo Research, #D4204). Before prepping, we recorded a 1.633 optical density.

### Electroporation

GM12878 cells were cultured so that cell density was between 400,000 and 800,000 cells/mL on day of transfection. Three replicates were performed on separate days. For each replicate, a total of 20 electroporation reactions was performed using the Neon™ Transfection System 100 µL Kit (Invitrogen, #MPK10025) and the associated Neon™ Transfection System (Invitrogen, #MPK5000). 121 million GM12878 cells were collected, washed with 45mL PBS, and resuspended in 2178μL Buffer R. For each reaction, 5 million cells (in 90μL Buffer R) were electroporated with 5μg of ATAC-STARR-seq plasmid DNA (in 10μL nuclease-free water) in a total volume of 100μL with the following parameters: 1100V, 30ms, and 2 pulses. Electroporated cells were dispensed immediately into a pre-warmed T-75 flask containing 50mL of RPMI 1640 with 20% fetal bovine serum and 2mM GlutaMAX.

### Cell Harvest

24 hours after transfection, the 50mL ATAC-STARR-seq flask was divided into two equal volumes; plasmid DNA was extracted from one volume, while reporter RNAs were extracted from the other. Plasmid DNA was isolated with the ZymoPURE II Plasmid Midiprep kit (#D4200) and eluted in 50μL 10mM Tris-HCL pH 8.0. Prior to lysis, cells were washed with 25mL PBS to remove any extracellular plasmid DNA. Reporter RNAs were extracted in a stepwise process. First, total RNA was isolated from the second volume of transfected cells using the TRIzol™ Reagent and Phasemaker™ Tubes Complete System (Invitrogen™, #A33251). Specifically, 5mL TRIzol was added to homogenize the washed and pelleted cells. Next, polyadenylated RNA was isolated from total RNA using oligo d(T)25 Magnetic Beads (NEB, #S1419S) at a 1μg Total RNA to 10μg beads ratio. We performed this step at 4°C and eluted into 50μL10mM Tris-HCl pH 7.5. The extracted poly(A)+ RNA was treated with DNase I (NEB, #M0303S). This reaction was cleaned up using the Zymo Research RNA Clean & Concentrator-25 kit (Zymo Research, #R1018).

### First-strand cDNA synthesis

For each sample, ten 50μL reverse transcription reactions were carried out using PrimeScript™ Reverse Transcriptase (Takara, #2680) and a gene specific primer (STARR_GSP) as described by Muerdter *et al*. 2018 (Muerdter et al. 2018). Single-stranded cDNA was treated with RNAse A at a concentration of 20μg/mL in low salt concentrations and cleaned up with a Zymo Research DNA Clean & Concentrator-5 kit.

### Illumina Sequencing Library Preparation

For reisolated plasmid and reporter RNA samples, Illumina-compatible libraries were generated using NEBNext^®^ Ultra™ II Q5^®^ Master Mix and a unique combination of the following Nextera indexes: N504-N505 (i5) and N701-N702 (i7), see Supplementary Table 1 for primer sequences. DNA samples were amplified for 8 PCR cycles, while RNA was amplified for 12-13 cycles. In both cases, products were purified with the Zymo Research DNA Clean & Concentrator-5 kit and analyzed for concentration and size distribution using a HSD5000 screentape. Purified products were sequenced on an Illumina NovaSeq, PE150, at a requested read depth of 50 or 75 million reads, for DNA and RNA samples, respectively, through the Vanderbilt Technology for Advanced Genomics (VANTAGE) sequencing core. Reads were processed and analyzed as described in the supplemental methods.

## DATA ACCESS

Raw data and processed results are available on Gene Expression Omnibus (GEO), accession number GSE181317 (private token: udibokwkjlazlmv). Python scripts and additional code for ATAC-STARR-seq data analysis are posted in a repository on the Hodges Lab GitHub (https://github.com/HodgesGenomicsLab/ATAC-STARR-seq) (Hansen and Hodges 2021). An interactive version of the protocol is posted on protocols.io (dx.doi.org/10.17504/protocols.io.b2nuqdew) and a pdf of the protocol at publication date is included as a supplementary data file.

## ACKNOWLEDGEMENTS

We thank Sarah Fong, Tony Capra, and members of the Hodges Lab, especially Kelly Barnett, Tim Scott, Lindsey Guerin, Verda Agan, Elizabeth Dorans, and Ali Wilt for helpful feedback and discussions. We also thank Biorender.com for illustrations, Addgene for plasmids used in the study, the Dave Cortez Lab for use of their Bio-Rad Gene Pulser, and the Manny Ascano Lab for qPCR primers and helpful advice. We are grateful for support of the project and the time invested in producing this manuscript by the NIH awards [K22 CA184308-03 to E.H], Department of Defense Idea Award [W81XWH-20-1-0522 to E.H], and American Cancer Society (ACS) Institutional Research Grant (#IRG-15-169-56).

## REFERENCES

Arnold CD, Gerlach D, Stelzer C, Boryn LM, Rath M, Stark A. 2013. Genome-wide quantitative enhancer activity maps identified by STARR-seq. Science 339: 1074–1077.

Barnett KR, Decato BE, Scott TJ, Hansen TJ, Chen B, Attalla J, Smith AD, Hodges E. 2020. ATAC-Me Captures Prolonged DNA Methylation of Dynamic Chromatin Accessibility Loci during Cell Fate Transitions. Mol Cell 77: 1350–1364 e1356.

Bentsen M, Goymann P, Schultheis H, Klee K, Petrova A, Wiegandt R, Fust A, Preussner J, Kuenne C, Braun T et al. 2020. ATAC-seq footprinting unravels kinetics of transcription factor binding during zygotic genome activation. Nat Commun 11: 4267.

Buenrostro JD, Giresi PG, Zaba LC, Chang HY, Greenleaf WJ. 2013. Transposition of native chromatin for fast and sensitive epigenomic profiling of open chromatin, DNA-binding proteins and nucleosome position. Nat Methods 10: 1213–1218.

Cao Z, Sun X, Icli B, Wara AK, Feinberg MW. 2010. Role of Kruppel-like factors in leukocyte development, function, and disease. Blood 116: 4404–4414.

Chaudhri VK, Dienger-Stambaugh K, Wu Z, Shrestha M, Singh H. 2020. Charting the cis-regulome of activated B cells by coupling structural and functional genomics. Nat Immunol 21: 210–220.

Consortium EP, Moore JE, Purcaro MJ, Pratt HE, Epstein CB, Shoresh N, Adrian J, Kawli T, Davis CA, Dobin A et al. 2020. Expanded encyclopaedias of DNA elements in the human and mouse genomes. Nature 583: 699–710.

Corces MR, Trevino AE, Hamilton EG, Greenside PG, Sinnott-Armstrong NA, Vesuna S, Satpathy AT, Rubin AJ, Montine KS, Wu B et al. 2017. An improved ATAC-seq protocol reduces background and enables interrogation of frozen tissues. Nat Methods 14: 959–962.

Dogan N, Wu W, Morrissey CS, Chen KB, Stonestrom A, Long M, Keller CA, Cheng Y, Jain D, Visel A et al. 2015. Occupancy by key transcription factors is a more accurate predictor of enhancer activity than histone modifications or chromatin accessibility. Epigenetics Chromatin 8: 16.

Doni Jayavelu N, Jajodia A, Mishra A, Hawkins RD. 2020. Candidate silencer elements for the human and mouse genomes. Nat Commun 11: 1061.

Gasperini M, Tome JM, Shendure J. 2020. Towards a comprehensive catalogue of validated and target-linked human enhancers. Nat Rev Genet 21: 292–310.

Glaser LV, Steiger M, Fuchs A, van Bommel A, Einfeldt E, Chung HR, Vingron M, Meijsing SH. 2021. Assessing genome-wide dynamic changes in enhancer activity during early mESC differentiation by FAIRE-STARR-seq. Nucleic Acids Res 49: 12178–12195.

Haberle V, Stark A. 2018. Eukaryotic core promoters and the functional basis of transcription initiation. Nat Rev Mol Cell Biol 19: 621–637.

Hansen TJ, Hodges E. 2021. Simultaneous profiling of regulatory activity, chromatin accessibility, and transcription factor occupancy with ATAC-STARR-seq (V1.0.0). githubcom/HodgesGenomicsLab/ATAC-STARR-seq doi:10.5281/zenodo.5764666.

Heinz S, Romanoski CE, Benner C, Glass CK. 2015. The selection and function of cell type-specific enhancers. Nat Rev Mol Cell Biol 16: 144–154.

Inoue F, Kircher M, Martin B, Cooper GM, Witten DM, McManus MT, Ahituv N, Shendure J. 2017. A systematic comparison reveals substantial differences in chromosomal versus episomal encoding of enhancer activity. Genome Res 27: 38–52.

Johnson GD, Barrera A, McDowell IC, D’Ippolito AM, Majoros WH, Vockley CM, Wang X, Allen AS, Reddy TE. 2018. Human genome-wide measurement of drug-responsive regulatory activity. Nat Commun 9: 5317.

Kim YS, Johnson GD, Seo J, Barrera A, Cowart TN, Majoros WH, Ochoa A, Allen AS, Reddy TE. 2021. Correcting signal biases and detecting regulatory elements in STARR-seq data. Genome Res doi:10.1101/gr.269209.120.

Kircher M, Xiong C, Martin B, Schubach M, Inoue F, Bell RJA, Costello JF, Shendure J, Ahituv N. 2019. Saturation mutagenesis of twenty disease-associated regulatory elements at single base-pair resolution. Nat Commun 10: 3583.

Klein JC, Agarwal V, Inoue F, Keith A, Martin B, Kircher M, Ahituv N, Shendure J. 2020. A systematic evaluation of the design and context dependencies of massively parallel reporter assays. Nat Methods 17: 1083–1091.

Klemm SL, Shipony Z, Greenleaf WJ. 2019. Chromatin accessibility and the regulatory epigenome. Nat Rev Genet 20: 207–220.

Lambert SA, Jolma A, Campitelli LF, Das PK, Yin Y, Albu M, Chen X, Taipale J, Hughes TR, Weirauch MT. 2018. The Human Transcription Factors. Cell 172: 650–665.

Lee D, Shi M, Moran J, Wall M, Zhang J, Liu J, Fitzgerald D, Kyono Y, Ma L, White KP et al. 2020. STARRPeaker: uniform processing and accurate identification of STARR-seq active regions. Genome Biol 21: 298.

Love MI, Huber W, Anders S. 2014. Moderated estimation of fold change and dispersion for RNA-seq data with DESeq2. Genome Biol 15: 550.

Maricque BB, Dougherty JD, Cohen BA. 2017. A genome-integrated massively parallel reporter assay reveals DNA sequence determinants of cis-regulatory activity in neural cells. Nucleic Acids Res 45: e16.

Melnikov A, Murugan A, Zhang X, Tesileanu T, Wang L, Rogov P, Feizi S, Gnirke A, Callan CG, Jr., Kinney JB et al. 2012. Systematic dissection and optimization of inducible enhancers in human cells using a massively parallel reporter assay. Nat Biotechnol 30: 271–277.

Muerdter F, Boryn LM, Woodfin AR, Neumayr C, Rath M, Zabidi MA, Pagani M, Haberle V, Kazmar T, Catarino RR et al. 2018. Resolving systematic errors in widely used enhancer activity assays in human cells. Nat Methods 15: 141–149.

Neumayr C, Pagani M, Stark A, Arnold CD. 2019. STARR-seq and UMI-STARR-seq: Assessing Enhancer Activities for Genome-Wide-, High-, and Low-Complexity Candidate Libraries. Curr Protoc Mol Biol 128: e105.

Pang B, Snyder MP. 2020. Systematic identification of silencers in human cells. Nat Genet 52: 254–263.

Patwardhan RP, Hiatt JB, Witten DM, Kim MJ, Smith RP, May D, Lee C, Andrie JM, Lee SI, Cooper GM et al. 2012. Massively parallel functional dissection of mammalian enhancers in vivo. Nat Biotechnol 30: 265–270.

Picelli S, Bjorklund AK, Reinius B, Sagasser S, Winberg G, Sandberg R. 2014. Tn5 transposase and tagmentation procedures for massively scaled sequencing projects. Genome Res 24: 2033–2040.

Roadmap Epigenomics C, Kundaje A, Meuleman W, Ernst J, Bilenky M, Yen A, Heravi-Moussavi A, Kheradpour P, Zhang Z, Wang J et al. 2015. Integrative analysis of 111 reference human epigenomes. Nature 518: 317–330.

Ryan GE, Farley EK. 2020. Functional genomic approaches to elucidate the role of enhancers during development. Wiley Interdiscip Rev Syst Biol Med 12: e1467.

Sambrook J, Russell DW. 2006. Standard ethanol precipitation of DNA in microcentrifuge tubes. CSH Protoc 2006.

Santiago-Algarra D, Dao LTM, Pradel L, Espana A, Spicuglia S. 2017. Recent advances in high-throughput approaches to dissect enhancer function. F1000Res 6: 939.

Thurman RE, Rynes E, Humbert R, Vierstra J, Maurano MT, Haugen E, Sheffield NC, Stergachis AB, Wang H, Vernot B et al. 2012. The accessible chromatin landscape of the human genome. Nature 489: 75–82.

Uhlen M, Fagerberg L, Hallstrom BM, Lindskog C, Oksvold P, Mardinoglu A, Sivertsson A, Kampf C, Sjostedt E, Asplund A et al. 2015. Tissue-based map of the human proteome. Science 347: 1260419.

Uhlen M, Karlsson MJ, Zhong W, Tebani A, Pou C, Mikes J, Lakshmikanth T, Forsstrom B, Edfors F, Odeberg J et al. 2019. A genome-wide transcriptomic analysis of protein-coding genes in human blood cells. Science 366.

Vierstra J, Lazar J, Sandstrom R, Halow J, Lee K, Bates D, Diegel M, Dunn D, Neri F, Haugen E et al. 2020. Global reference mapping of human transcription factor footprints. Nature 583: 729–736.

Wang X, He L, Goggin SM, Saadat A, Wang L, Sinnott-Armstrong N, Claussnitzer M, Kellis M. 2018. High-resolution genome-wide functional dissection of transcriptional regulatory regions and nucleotides in human. Nat Commun 9: 5380.

Yan F, Powell DR, Curtis DJ, Wong NC. 2020. From reads to insight: a hitchhiker’s guide to ATAC-seq data analysis. Genome Biol 21: 22.

