## Supplemental Material for "Identifying transcription factor-bound activators and silencers in the chromatin accessible human genome using ATAC-STARR-seq"

**<u>Supplemental Materials for:</u>**

**<u>Table of Contents:</u>**

**Supplemental Text**

ATAC-STARR-seq Plasmid library complexity……….4

Optimizing ATAC-STARR-seq assay timeframe……….4

Investigating the influence of replicates on region calls……….5

Duplicate removal hinders region calling sensitivity………6

**Supplemental Methods**

Determination of Harvest Time with Quantitative PCR……….6

Plasmid Library Complexity Estimation……….7

Transfection efficiency Estimation……….8

Read Processing……….8

Accessibility Analysis……….9

Active and Silent Region Calling……….10

Replicate Count Effects……….11

Orientation Analysis……….12

Active and Silent Peak Characterization……….13

TF Footprinting……….14

Multiomic Integration……….15

**Supplemental References**

**Supplemental Tables**

ST1. ATAC-STARR-seq Sequencing Summary Statistics……….18

ST2. Genrich peak counts for varying FDR thresholds……….18

**Supplemental Figures**

S1. ATAC-STARR Optimization……….19

S2. Characterization of ATAC-STARR sequencing Libraries……….20

S3. Correlation between ATAC-STARR-seq replicates……….21

S4. Comparison between the sliding window and the fragment group active region calling methods……….22

S5. Analysis of replicate count on region calling sensitivity……….23

S6. Comparison between keeping duplicates and removing duplicates to call active regions……….24

S7. Assessment of potential orientation bias in ATAC-STARR-seq data……….25

S8. Histone modification ChIP-seq signal at accessible chromatin peaks……….26

### SUPPLEMENTAL TEXT

#### ATAC-STARR-seq plasmid library complexity

A successful ATAC-STARR-seq experiment is predicated on maintaining complexity at all stages of the protocol. We estimated the initial complexity of our ATAC-STARR-seq plasmid library by sequencing the library at low depth and estimating the number of unique reads with the Preseq software package (Daley and Smith 2013) (Supplementary Figure 1A). The GM12878 ATAC-STARR-seq plasmid library contains a maximum complexity of about 50 million unique accessible DNA fragments, providing ample coverage of accessible loci.

#### Optimizing ATAC-STARR-seq assay timeframe

The introduction of plasmid DNA into cells produces an interferon-stimulated gene response that can confound the isolation of biologically relevant regulatory activity (Muerdter et al. 2018). To minimize this interference in our data, we determined the optimal incubation time between electroporation and harvest. Two factors play an important role in determining when to harvest RNA: global reporter RNA expression levels and the timing of interferon stimulated gene response to STARR-seq reporter plasmid DNA. To investigate both factors, we electroporated ATAC-STARR-seq plasmid DNA, isolated poly-adenylated RNA at several time points after transfection, quantified RNA expression with qPCR, and compared to an untransfected sample (Supplementary Figure 1A). An increase in reporter RNA expression is observed at 3 hours (the earliest timepoint) and remains stable at later time points. We measured expression of IFNB1, IFIT2, and ISG15 to characterize the interferon stimulated gene response in our system. RNA expression for all three genes increases initially but returns to baseline by 24 hours. Given the persistent level of reporter RNAs and the attenuated interferon stimulated gene response in our system, we decided to harvest 24 hours after electroporation. Together, this allows us to capture reporter RNAs that reflect steady-state regulatory properties of GM12878 accessible regions without sacrificing reporter RNA recovery.

**Investigating the influence of replicates on region calls**

We note that the Wang *et al*. 2018 study reported twice the number of active regions reported herein. This discrepancy may be explained in part by the use of the super core promoter in their assay, but another major difference between the two studies is replicate number (five replicates versus three replicates). To determine if the difference in active region count is driven by replicate number, we downloaded and analyzed raw sequencing data from Wang *et al.* 2018 using our pipeline and analysis methods. We then assigned reads to the bins we analyzed and called active regions using either three or five replicates (Supplementary Figure 5A). With five replicates, we also captured ~66,000 active regions; however, we identified ~39,000 regions with only three replicates. This is much closer to the number we report (~30,000) and suspect the extra 9,000 regions may be the result of experimental differences, such as the promoter employed. Altogether the number of called active regions increases with more replicates.

To further investigate the effect of replicate number on region calling sensitivity in our data, we merged and split our three ATAC-STARR-seq replicates into five randomly sampled “pseudo-replicates”. We then called active regions using two, three, four, or five pseudo-replicates (Supplementary Figure 5B). We find the largest increase in region count going from two to three replicates. Thus, the three replicate condition seems to yield the best value, while additional replicates may be needed to detect more weakly active regulatory regions. However, it is also very important to note that studies investigating the relationship between replicate number, sensitivity, and accuracy for RNA-seq data have demonstrated that performing more replicates yields more differentially expressed genes, but this is concomitant with an increase in false positive rate (Schurch et al. 2016; Lamarre et al. 2018). Therefore, the additional regions that are called with increasing replicate counts may represent a disproportionate number of false positives and may affect the outcomes of certain accuracy-sensitive applications like computational modelling.

**Duplicate removal hinders region calling sensitivity**

A question that often arises when determining biological signals from sequence read count data is whether to collapse read duplicates, as duplicates can arise both technically (PCR duplicates) and biologically (active regions generate multiple transcripts of themselves). To understand their contribution to data interpretation, we analysed our data with and without duplicates and compared the output. Removal of duplicates produces modest improvements to correlation coefficients between replicates, although both conditions had correlations indicative of satisfactory reproducibility (Supplementary Figures 3, 6A-B). However, excluding duplicates produced many fewer active regions called than including duplicates (~21,000 fewer regions) (Supplementary Figure 6C). Together, this indicates that removing duplicates modestly improves reproducibility but significantly sacrifices sensitivity. Furthermore, most of the regions called without duplicates are also called when duplicates are included, indicating that, for the most part, duplicate removal affects sensitivity and not accuracy (Supplementary Figure 6D). Because the with-duplicate analysis yielded many more additional regions and is reproducible between replicates, we included duplicates in our activity analysis moving forward. Importantly, because our approach filters by significance, reproducibility is required when calling active and silent regions. Therefore, identified active and silent regions are of high confidence when including duplicates.

### SUPPLEMENTAL METHODS

**Determination of Harvest Time with Quantitative PCR**

GM12878 cells were cultured so that cell density was between 400,000 and 800,000 cells/mL on day of transfection. Three replicates were performed on separate days. For each sample, 5 million GM12878 cells were electroporated with 5µg ATAC-STARR-seq plasmid DNA using the Neon™ Transfection System 100 µL Kit (Invitrogen, #MPK10025) and the associated Neon™ Transfection System (Invitrogen, #MPK5000) in Buffer R with the following parameters: 1100V, 30ms, and 2 pulses. Electroporated cells were dispensed immediately into pre-warmed T-12.5 flasks containing 6.25mL of RPMI 1640 with 20% fetal bovine serum and 2mM GlutaMAX.

Total RNA was harvested at various time points—3hr, 6hr, 12hr, 24hr, and 36hr—using the TRIzol™ Reagent and Phasemaker™ Tubes Complete System (Invitrogen™, #A33251). For each sample, 0.75mL TRIzol was added to cell pellets. First-strand cDNA synthesis was performed using an Oligo $(dT)_{25}$ primer and the SuperScript™ IV First-Strand Synthesis System (Invitrogen™, #18091050). cDNA was treated with RNase H to remove RNA from RNA-DNA dimers. For each replicate, 10µL quantitative PCR reactions were performed in technical triplicate using PowerUp™ SYBR™ Green Master Mix (Applied Biosystems™, #A25742) on a StepOnePlus™ Real-Time PCR System (Applied Biosystems™, #4376600). For each reaction, 1µL of the reverse-transcribed product was added and gene-specific primers were supplied at a final concentration of 500nM (see Supplementary Table 3 for primer sequences). Fold-change was calculated with the ΔΔCt method, using either GAPDH or ACTB as the housekeeping gene for reporter RNA or ISG targets, respectively. Plots were made with ggplot2 (version 3.3.5) (Wickham 2016) in R (version 4.1.1).

**Plasmid Library Complexity Estimation**

Plasmid inserts were amplified via PCR from 3.75µg ATAC-STARR-seq plasmid library using NEBNext® Ultra™ II Q5® Master Mix and the Nextera indexes, N505 and N701, see Supplementary Table 3 for primer sequences. Products were purified with the Zymo Research DNA Clean & Concentrator-5 kit

(#D4013) and analyzed for concentration and size distribution using a HSD5000 screentape. Purified products were sequenced on an Illumina NovaSeq, PE150, at a requested read depth of 25 million reads through the Vanderbilt Technology for Advanced Genomics (VANTAGE) sequencing core.

**Transfection efficiency estimation**

Transfection efficiency is a critical ATAC-STARR-seq bottleneck, particularly for difficult to transfect cells like GM12878. In parallel with ATAC-STARR-seq, we electroporated GM12878 cells with a pcDNA3.1-eGFP plasmid and estimated transfection efficiency as the percentage of GFP positive cells when measured by flow cytometry 24 hours later. Specifically, GM12878 cells were electroporated following same conditions as above with either purified pcDNA3.1-eGFP plasmid or nuclease-free water and then prepared for flow cytometry 24 hours later at a concentration of $1.25 \times 10^6$ cells/mL in 1xPBS solution containing 1% BSA. We halved both GFP and water samples and stained one half of each with propidium iodide (Sigma-Aldrich, #P4864). Unstained cells (water/PI-) were used in conjunction with compensation control cells (GFP/PI- or water/PI+) to quantify the percentage of living GFP positive cells in the experimental condition (GFP/PI+) via flow cytometry; this percentage was the reported transfection efficiency. When performed in parallel to ATAC-STARR-seq plasmid library transfection, we consistently achieve around 10-20% efficiency (data not shown).

**Read Processing**

FASTQ files for all four ATAC-seq replicates from Buenrostro *et al*. 2013 and all five HiDRA replicates from Wang *et al.* 2018 were downloaded from the NCBI sequence read archive (run codes: SRR891268-SRR891271 and SRR6050484-SRR6050523, respectively) and were processed using the same pipeline as ATAC-STARR-seq (Buenrostro et al. 2013; Wang et al. 2018). For this publicly available data and our own, FASTQ files were trimmed and analysed for quality with Trim Galore! (version 0.6.7, https://www.bioinformatics.babraham.ac.uk/projects/trim_galore) using the --fastqc and --paired parameters. Trimmed reads were mapped to hg38 with bowtie2 (version 2.3.5.1) using the following parameters: -X 500 --sensitive --no-discordant --no-mixed (Langmead and Salzberg 2012). Mapped reads were filtered to remove reads with MAPQ < 30, reads mapping to mitochondrial DNA, and reads mapping to ENCODE blacklist regions using a variety of functions from the Samtools software package (version 1.13) (Li et al. 2009). When desired, duplicates were removed with the *markDuplicates* function from Picard (version 2.26.3) (https://broadinstitute.github.io/picard/). Read count was determined using the *flagstat* function from Samtools. Read counts for each step are provided in Supplementary Table 1. We also provide a python script on our GitHub repository (Hansen and Hodges 2021) that performs the processing steps above. Complexity was estimated using the *lc-extrap* function from the Preseq package (version 2.0.0) (Daley and Smith 2013) and insert size was determined using the *CollectInsertSizeMetrics* function from Picard. Complexity curves were plotted in R with ggplot2.

**Accessibility Analysis**

*Peak Calling.* We called accessibility peaks with the Genrich software package (version 0.5, https://github.com/jsh58/Genrich), using deduplicated bam files. For ATAC-STARR-seq, we used all three replicates of reisolated plasmid samples. For Buenrostro data, we used all four of the available replicates. For both, we set a false-discovery rate of 0.0001 and the -j parameter, which specifies ATAC-seq mode.

*Peak Comparisons*. Peaks between Buenrostro and ATAC-STARR-seq plasmid DNA were compared using the *jaccard* function from the BEDtools package (version 2.30.0) (Quinlan and Hall 2010). FRiP scores (the fraction of reads in peaks) and the genomic fraction represented by each peak set was calculated using custom code available on our GitHub repository. Euler plots were made in R with the eulerr package (version 6.1.0) (Larsson 2021) and bar charts were made in R with ggplot2.

*Signal Tracks.* Accessibility signal tracks were generated with the *bamCoverage* function from the deepTools package (version 3.5.1) (Ramirez et al. 2016) using the following parameters: -bs 10 --normalizeUsing CPM -e --centerReads. Signal was plotted using the Sushi package (version 1. 30.0) (Phanstiel et al. 2014) in R.

**Active and Silent Region Calling**

We called active and silent regions using the sliding window and fragment groups methods. In both cases, except where specified, mapped read files containing duplicates were used for region calling. Overlap between the two active region sets identified by each method was determined using BEDtools jaccard. Methods for each are listed below.

*Sliding Window.* Within ATAC-STARR-defined open chromatin regions, we generated 50 bp genomic, sliding window bins with a 10bp step size using the *makewindows* function and -s 10 -w 50 parameters from the BEDtools software package. Bins smaller than 50bp were removed from the analysis and reads were counted per bin for each replicate using the *featureCounts* function from the Subread package with the following parameters: -p -B -O --minOverlap 1 (Liao et al. 2014). The resulting counts matrix was pre-filtered to remove bins with zero counts and then analyzed with the DESeq2 software package (version 1.32.0) in R to identify active and silent bins (Love et al. 2014). Bins with an Benjamini–Hochberg (BH) adjusted p-value < 0.1 and $\log_2$ fold-change (RNA/DNA) > 0 were defined as active, whereas silent had a BH adjusted p-value < 0.1 and $\log_2$ fold-change (RNA/DNA) < 0. Overlapping and book-ended bins were merged with the *merge* function from BEDtools (using default parameters), resulting in active and silent regions. A python script for region calling is available on our GitHub repository. For the sliding window strategy, we also performed the analysis with or without duplicates in order to compare the results. For the without-duplicate analysis, deduplicated bam files were used at the *featureCounts* step, otherwise all parameters were the same. Active regions were compared using the *jaccard* function from the BEDtools package. Scatter plots and correlation coefficients for replicate-to-replicate comparisons were generated by first extracting DESeq-normalized counts, using the *counts(normalized=TRUE)* function, plotted using ggplot2, and compared using the *cor.test()* function in R using both spearman and pearson correlation methods.

*Fragment Groups.* We generated fragment groups using custom code based on the method described in Wang *et al* 2018 (Wang et al. 2018). Paired-end mapped reads were converted from bam to bed format using the *bamtobed* function from the BEDtools software package with option -bedpe and a custom *awk* function. Overlapping paired-end fragments were grouped using the *bedmap* function from the BEDOPS software package (version 2.4.28) (Neph et al. 2012) using the following parameters: --count --echo-map-range --fraction-both 0.75. Importantly, only fragment groups made up of 10 or more reads were used for downstream analysis. Reads were counted per fragment group for each replicate bam file using the *featureCounts* function from the Subread package (version 2.0.1) with the following parameters: -p -B -O --minOverlap 1. The resulting counts matrix was pre-filtered to remove bins with zero counts and then analyzed with the DESeq2 software package in R to identify active fragment groups. Fragment groups with an adjusted p-value < 0.1 and $log_2$ fold-change (RNA/DNA) > 0 were defined as active. This method resulted in many fragment groups that overlapped each other, so we isolated the most active region within each overlap using a custom function available on our GitHub repository; the resulting, non-redundant regions were defined as active peaks.

**Replicate Count Effects**

*HiDRA replicate count comparison.* Raw HiDRA sequencing data was downloaded and processed as described in the read processing section above. Using the same bins generated and analyzed in the active and silent region calling section, reads from all five HiDRA replicates were counted per bin using the *featureCounts* function from the Subread package and the following parameters: -p -B -O --

minOverlap 1. Active and regions were called in the same manner as described in the active and silent region calling section using either three or five replicates. Region counts for each condition were plotted using ggplot2.

*Pseudo-replicate analysis.* To create pseudo-replicates, all three replicate bam files of our ATAC-STARR data were merged using Samtools *merge.* Merged reads were split into five separate files using the Samtools *view* command with the *-s* options set to $rep.2, where .2 represents 20% of the reads and $rep represents the seed number for random sampling. In this way, each pseudo-replicate was sampled with a unique seed number and should, therefore differ from the other pseudo-replicates. Using the same bins analyzed in the active and silent region calling section, reads from all five pseudo-replicates were counted per bin and active regions were called in the same manner as described in the active and silent region calling section using two, three, four, or five pseudo-replicates. Region counts for each condition were plotted using ggplot2.

**Orientation Analysis**

Replicate bam files were merged using Samtools *merge*. Reads were split by orientation using Samtools *view -f*, which selects reads based on their SAM flags. Reads with flags 99 and 147 were assigned to the 5'-3' bam file, while reads with flags 83 and 163 were assigned to the 3'-5' bam file. The same bins generated and analyzed for region calling were used. Bins designated as active and silent were used for the active only and silent only analysis, respectively. The three bin sets were further subset into proximal and distal based on distance to the nearest TSS using the ChIPSeeker software package; proximal bins were defined as 2kb upstream and 1kb downstream of a TSS while distal was everything else. For each subset of bins, reads were counted per bin for the orientation-specific bam files using the *featureCounts* function from the Subread package with the following parameters: -p -B -O --minOverlap 1. Scatter plots of counts per million normalize read count were generated with ggplot2 and both spearman and pearson correlation coefficients were determined with the *cor.test()* function in R. Bins with a greater than 5 read count difference between insert orientations were considered to be biased; we based this threshold on the all distal bins scatterplot with the assumption that distal bins should not display an orientation bias. The percentage biased was plotted with ggplot2.

**Active and Silent Peak Characterization**

*Annotation.* Active and silent peak sets were annotated relative to transcription start site (TSS) locations and plotted in R using the ChIPSeeker package (version 1.28.3) (Yu et al. 2015); promoters were defined as 2kb upstream and 1kb downstream of a TSS. ChromHMM state was assigned to each peak using the BEDtools *intersect* function and -u parameter; the list of hg38 18-state ChromHMM regions (Roadmap Epigenomics et al. 2015) (https://egg2.wustl.edu/roadmap/data/byFileType/chromhmmSegmentations/ChmmModels/core_K27ac/jointModel/final/E116_18_core_K27ac_hg38lift_mnemonics.bed.gz) were intersected against the regions sets of interest and the proportion was plotted with ggplot2.

*Heatmaps.* The activity bigwig was generated with the deepTools package. Merged bam files for RNA and DNA were converted to counts per million normalized bedGraph files using the *bamCoverage* function and the following parameters: -bs 10 --normalizeUsing CPM. The resulting RNA bigwig was normalized to the DNA bigwig to generate a signal file of log2(RNA/DNA) ratio using the *bigwigCompare* function and the following parameters: -bs 1 --operation log2 --pseudocount 1 –skipZeroOverZero. Heatmaps were generated using the deepTools package. Activity signal was plotted at distal and proximal regions and region order was ranked by maximum mean signal. GM12878 ChIP-seq bigwig files were downloaded from the ENCODE consortium (Consortium et al. 2020) and plotted. The matrix was made using the *computeMatrix* function, with the following parameters: -a 2000 -b 2000 --

referencePoint center -bs 10 --missingDataAsZero. The matrix was plotted using the *plotHeatmap* function with the following key parameters: --sortUsing mean --sortUsingSamples 1.

*Histone Signal Boxplots.* We intersected silent and active regions with our accessible peaks file using the *intersect* function from the BEDtools software package to get peaks that contain an active region, a silent region, both an active and silent region, or neither. Using the *slop* function from BEDtools we then extended ChrAcc peaks by 1kb on either side and then used the bigwigCompare function from the DeepTools package to determine H3K4me1/H3K4me3/H3K27ac/H3Kme3 GM12878 ChIP-seq bigwig signal distributions for each for the ChrAcc peak types. The same ENCODE files used in the heatmap analysis above, were also used here. The plotted values represent the average *fold-change over control* for each ChrAcc peak +/- 1kb. Plots were made with ggplot2.

*Motif enrichment.* We performed motif enrichment on the active and silent peak sets using the findMotiftsGenome.pl script from the HOMER package (version 4.10, http://homer.ucsd.edu/) (Duttke et al. 2019) using the following parameters: -size given -mset vertebrates. Plots were made with ggplot2.

**TF footprinting**

*Computational footprinting.* Transcription factor footprinting was performed using the TOBIAS software package (version 0.12.12) (Bentsen et al. 2020). Deduplicated mapped reads were used to generate Tn5-bias corrected bigwig signal files using the *ATACorrect* function. Using the corrected signal files, TF binding was calculated with the *ScoreBigWig* function and footprints for individual TFs were called for all core non-redundant vertebrate JASPAR motifs (Fornes et al. 2020) using the *BINDetect* function. Motifs with a footprint were classified as "bound", while motifs without a footprint were classified as "unbound". The "archetype" for each TF was assigned by cross-referencing the motif annotations table from Viestra *et al*. 2020 (Vierstra et al. 2020).

*Data Visualization.* Heatmaps were generated using the deepTools package. GM12878 ChIP-seq bigwig files were downloaded from ENCODE (www.encodeproject.org) (Consortium et al. 2020) and plotted with Tn5-corrected signal at all accessible CTCF and CREB/ATF/1 motifs (defined as the "all" bed file for CTCF or JUNB from BINDetect) using the *computeMatrix reference-point* function with the following key parameters: -a 200 -b 200 --referencePoint center --missingDataAsZero -bs1. The resulting matrix was plotted using the *plotHeatmap* function and the following key parameters: --sortUsing mean --sortUsingSamples 1. Aggregate plots were also generated using the deepTools package. Tn5-corrected signal was measured at bound and unbound sites for each TF archetype using the *computeMatrix reference-point* function with the following key parameters: -a 75 -b 75 --referencePoint center --missingDataAsZero -bs 1. The resulting matrix was plotted using the *plotProfile* function.

**Multiomic Integration**

Signal and regions were visualized at the given locus using the Sushi package in R. To determine the presence or absence of a TF footprint, we intersected TF footprints with the active and silent regions bed file and reported +/- for presence of the footprint using custom code available on our GitHub repository. Footprints were selected based on top hits from the motif enrichment analysis above. Active and silent regions without a footprint for the queried TFs were removed from the analysis. We clustered the region subsets with the pheatmap package (version 1.0.12, https://github.com/raivokolde/pheatmap), using the clustering_distance_row/columns = "binary" parameter; we cut the tree into 6 clusters for active and silent. We extracted the regions from each cluster and then, using the ChIPSeeker package, assigned the nearest neighbor gene. Using ClusterProfiler (Yu et al. 2012) and ReactomePA (Jassal et al. 2020), we then performed reactome pathway enrichment analysis on the nearest neighbor gene sets. We applied a 0.05 and 0.1 p-value cut-off for active and silent clusters, respectively.

### SUPPLEMENTAL REFERENCES


Bentsen M, Goymann P, Schultheis H, Klee K, Petrova A, Wiegandt R, Fust A, Preussner J, Kuenne C, Braun T et al. 2020. ATAC-seq footprinting unravels kinetics of transcription factor binding during zygotic genome activation. *Nat Commun* **11**: 4267.

Buenrostro JD, Giresi PG, Zaba LC, Chang HY, Greenleaf WJ. 2013. Transposition of native chromatin for fast and sensitive epigenomic profiling of open chromatin, DNA-binding proteins and nucleosome position. *Nat Methods* **10**: 1213-1218.

Consortium EP, Moore JE, Purcaro MJ, Pratt HE, Epstein CB, Shoresh N, Adrian J, Kawli T, Davis CA, Dobin A et al. 2020. Expanded encyclopaedias of DNA elements in the human and mouse genomes. *Nature* **583**: 699-710.

Daley T, Smith AD. 2013. Predicting the molecular complexity of sequencing libraries. *Nat Methods* **10**: 325-327.

Duttke SH, Chang MW, Heinz S, Benner C. 2019. Identification and dynamic quantification of regulatory elements using total RNA. *Genome Res* **29**: 1836-1846.

Fornes O, Castro-Mondragon JA, Khan A, van der Lee R, Zhang X, Richmond PA, Modi BP, Correard S, Gheorghe M, Baranasic D et al. 2020. JASPAR 2020: update of the open-access database of transcription factor binding profiles. *Nucleic Acids Res* **48**: D87-D92.

Hansen TJ, Hodges E. 2021. Simultaneous profiling of regulatory activity, chromatin accessibility, and transcription factor occupancy with ATAC-STARR-seq (V1.0.0). *githubcom/HodgesGenomicsLab/ATAC-STARR-seq* doi:10.5281/zenodo.5764666.

Jassal B, Matthews L, Viteri G, Gong C, Lorente P, Fabregat A, Sidiropoulos K, Cook J, Gillespie M, Haw R et al. 2020. The reactome pathway knowledgebase. *Nucleic Acids Res* **48**: D498-D503.

Lamarre S, Frasse P, Zouine M, Labourdette D, Sainderichin E, Hu G, Le Berre-Anton V, Bouzayen M, Maza E. 2018. Optimization of an RNA-Seq Differential Gene Expression Analysis Depending on Biological Replicate Number and Library Size. *Front Plant Sci* **9**: 108.

Langmead B, Salzberg SL. 2012. Fast gapped-read alignment with Bowtie 2. *Nat Methods* **9**: 357-359.

Larsson J. 2021. eulerr: Area-Proportional Euler and Venn Diagrams with Ellipses.

Li H, Handsaker B, Wysoker A, Fennell T, Ruan J, Homer N, Marth G, Abecasis G, Durbin R, Genome Project Data Processing S. 2009. The Sequence Alignment/Map format and SAMtools. *Bioinformatics* **25**: 2078-2079.

Liao Y, Smyth GK, Shi W. 2014. featureCounts: an efficient general purpose program for assigning sequence reads to genomic features. *Bioinformatics* **30**: 923-930.

Love MI, Huber W, Anders S. 2014. Moderated estimation of fold change and dispersion for RNA-seq data with DESeq2. *Genome Biol* **15**: 550.

Muerdter F, Boryn LM, Woodfin AR, Neumayr C, Rath M, Zabidi MA, Pagani M, Haberle V, Kazmar T, Catarino RR et al. 2018. Resolving systematic errors in widely used enhancer activity assays in human cells. *Nat Methods* **15**: 141-149.

Neph S, Kuehn MS, Reynolds AP, Haugen E, Thurman RE, Johnson AK, Rynes E, Maurano MT, Vierstra J, Thomas S et al. 2012. BEDOPS: high-performance genomic feature operations. *Bioinformatics* **28**: 1919-1920.

Phanstiel DH, Boyle AP, Araya CL, Snyder MP. 2014. Sushi.R: flexible, quantitative and integrative genomic visualizations for publication-quality multi-panel figures. *Bioinformatics* **30**: 2808-2810.

Quinlan AR, Hall IM. 2010. BEDTools: a flexible suite of utilities for comparing genomic features. *Bioinformatics* **26**: 841-842.

Ramirez F, Ryan DP, Gruning B, Bhardwaj V, Kilpert F, Richter AS, Heyne S, Dundar F, Manke T. 2016. deepTools2: a next generation web server for deep-sequencing data analysis. *Nucleic Acids Res* **44**: W160-165.

Roadmap Epigenomics C, Kundaje A, Meuleman W, Ernst J, Bilenky M, Yen A, Heravi-Moussavi A, Kheradpour P, Zhang Z, Wang J et al. 2015. Integrative analysis of 111 reference human epigenomes. *Nature* **518**: 317-330.

Schurch NJ, Schofield P, Gierlinski M, Cole C, Sherstnev A, Singh V, Wrobel N, Gharbi K, Simpson GG, Owen-Hughes T et al. 2016. How many biological replicates are needed in an RNA-seq experiment and which differential expression tool should you use? *RNA* **22**: 839-851.

Vierstra J, Lazar J, Sandstrom R, Halow J, Lee K, Bates D, Diegel M, Dunn D, Neri F, Haugen E et al. 2020. Global reference mapping of human transcription factor footprints. *Nature* **583**: 729-736.

Wang X, He L, Goggin SM, Saadat A, Wang L, Sinnott-Armstrong N, Claussnitzer M, Kellis M. 2018. High-resolution genome-wide functional dissection of transcriptional regulatory regions and nucleotides in human. *Nat Commun* **9**: 5380.

Wickham H. 2016. ggplot2: Elegant Graphics for Data Analysis.

Yu G, Wang LG, Han Y, He QY. 2012. clusterProfiler: an R package for comparing biological themes among gene clusters. *OMICS* **16**: 284-287.

Yu G, Wang LG, He QY. 2015. ChIPseeker: an R/Bioconductor package for ChIP peak annotation, comparison and visualization. *Bioinformatics* **31**: 2382-2383.

### SUPPLEMENTARY TABLES

#### Supplementary Table 1. ATAC-STARR-seq Sequencing Summary Statistics.

| Metric | DNA Rep 1 | DNA Rep 2 | DNA Rep 3 | RNA Rep 1 | RNA Rep 2 | RNA Rep 3 |
|---|---|---|---|---|---|---|
| ***Total read count (paired end)*** | 55,453,364 | 47,609,989 | 81,350,911 | 101,163,327 | 122,274,760 | 103,410,392 |
| ***Filtered read count (paired end)*** | 30,803,098 | 26,530,451 | 44,046,983 | 56,307,716 | 67,956,476 | 56,098,454 |
| ***Filtered & deduplicated read count (paired end)*** | 22,626,181 | 20,015,687 | 28,369,114 | 11,385,851 | 8,122,462 | 9,285,796 |
| ***Trimming Rate*** | 76% | 79% | 82% | 76% | 76% | 76% |
| ***Mapping Rate (>30MAPQ)*** | 61% | 61% | 59% | 61% | 61% | 59% |
| ***% mtDNA reads*** | 8.6% | 8.7% | 8.6% | 8.6% | 8.6% | 8.3% |
| ***% ENCODE blacklist reads*** | 0.05% | 0.05% | 0.05% | 0.05% | 0.05% | 0.05% |
| ***Duplication rate*** | 27% | 25% | 35.6% | 80% | 88% | 83% |
| ***Number of PCR Cycles*** | 8 | 8 | 8 | 13 | 13 | 12 |
| ***FastQC fields failed*** | Per base sequence content, Sequence Duplication Levels | | | | | |

#### Supplementary Table 2. Genrich peak counts for varying FDR thresholds.

| Sample | FDR < 0.01 | FDR < 0.001 | FDR < 0.0001 | FDR < 0.00001 |
|---|---|---|---|---|
| ***Buenrostro*** | 196,382 | 111,588 | 82,337 | 65,581 |
| ***ATAC-STARR*** | 162,877 | 124,612 | 101,904 | 85,668 |

**Supplementary Table 3 contains oligo sequences used in ATAC-STARR-seq and qPCR. It is included as a separate excel file.**

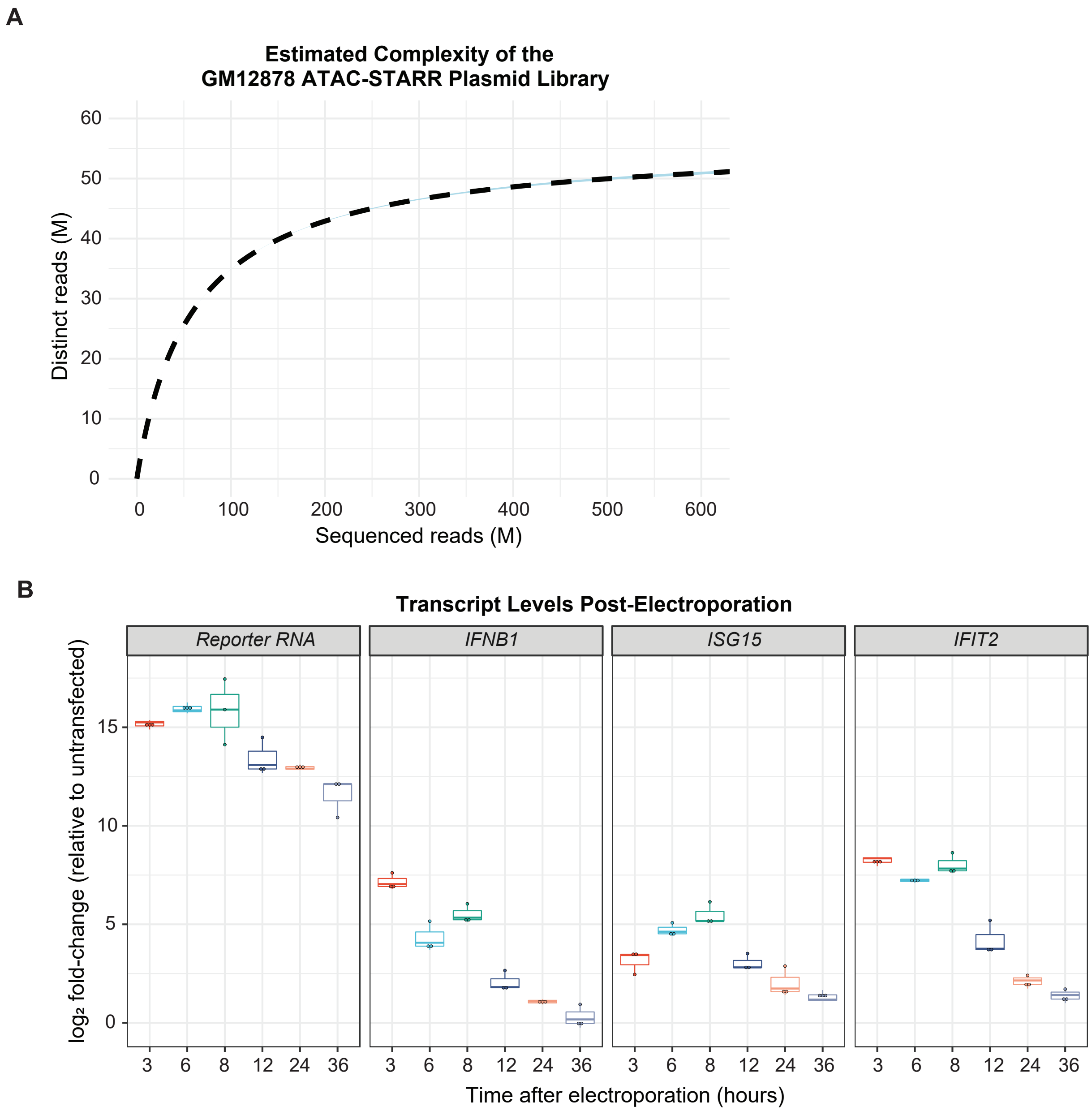


**Supplementary Figure 1. ATAC-STARR Optimization.** (A) Estimated complexity curve for the GM12878 ATAC-STARR plasmid library. Dashed lines represent predicted values from Preseq's lc-extrap. The associated ribbon plots (light blue) represent the 95% confidence interval reported with the predicted value. (B) Relative expression of reporter RNAs and three interferon-stimulated genes (IFNB1, IFIT2, and ISG15) at varying timepoints between 0- and 36-hours post-electroporation. For each analysis, fold-change values are relative to the untransfected condition. Three replicates were isolated and quantified for each timepoint.

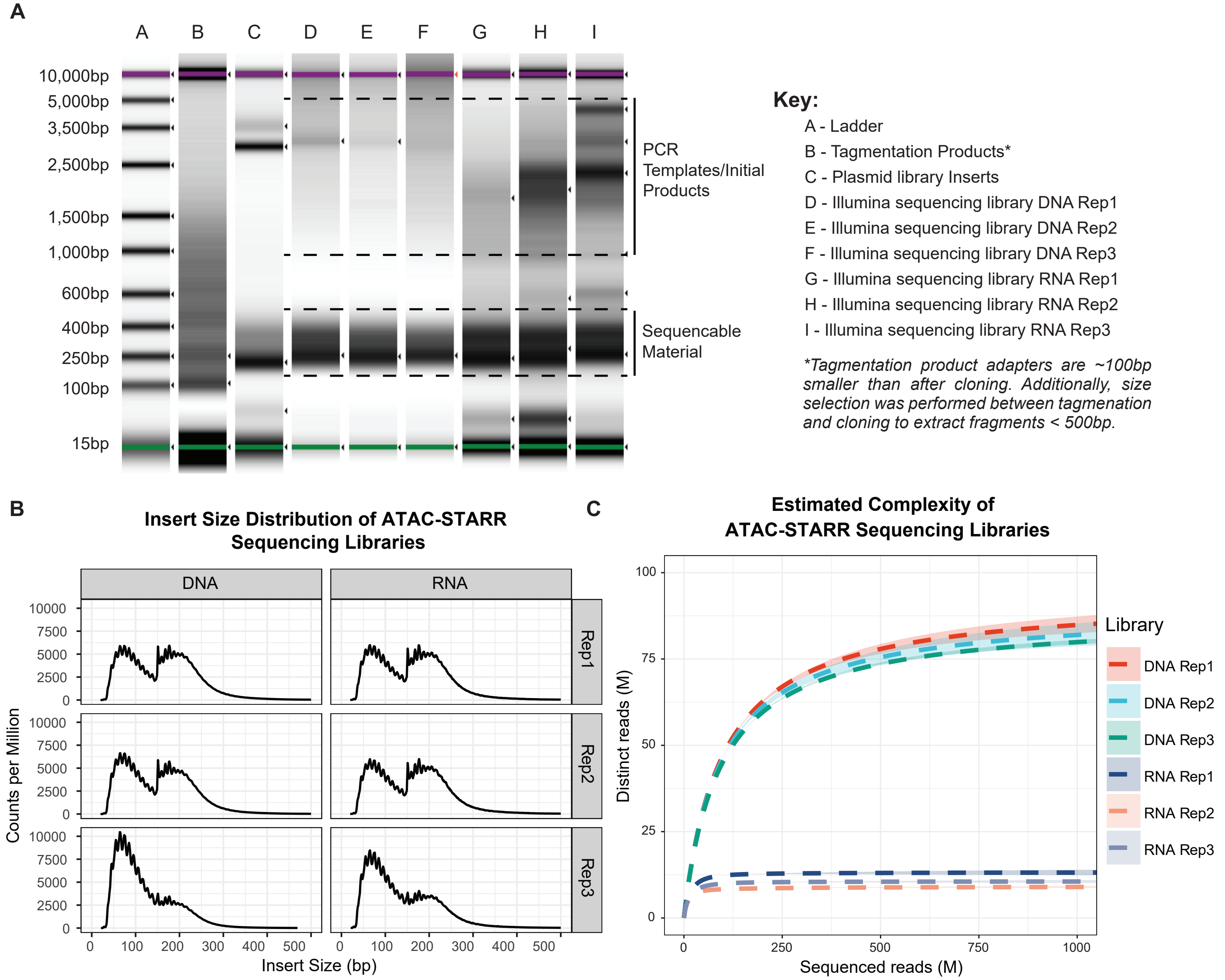


**Supplementary Figure 2. Characterization of ATAC-STARR sequencing libraries.** (A) Agilent Tapestation results for relevant steps of ATAC-STARR, this includes the following: tagmented products, plasmid library inserts, and illumina sequencing libraries for all three replicates of DNA and RNA. Tagmented products lack the full illumina adapter and therefore are about 100bp smaller than their later-stage counterparts. They also include larger fragments which were removed via selection before the cloning step. The Illumina-ready libraries were amplified using a minimal PCR cycle number and therefore the plasmid or cDNA template as well as the first and second round products can be seen as larger material—this material is not sequencable as it lacks at least one of the adapters required for cluster amplification. (B) Insert size distribution of ATAC-STARR-seq reads, as quantified by Picard's *CalculateInsertSizeMetrics*. (C) Estimated complexity curves for ATAC-STARR sequencing libraries. Dashed lines represent predicted values from Preseq's *lc-extrap*. The associated ribbon plots (light blue) represent the 95% confidence interval reported with the predicted value.

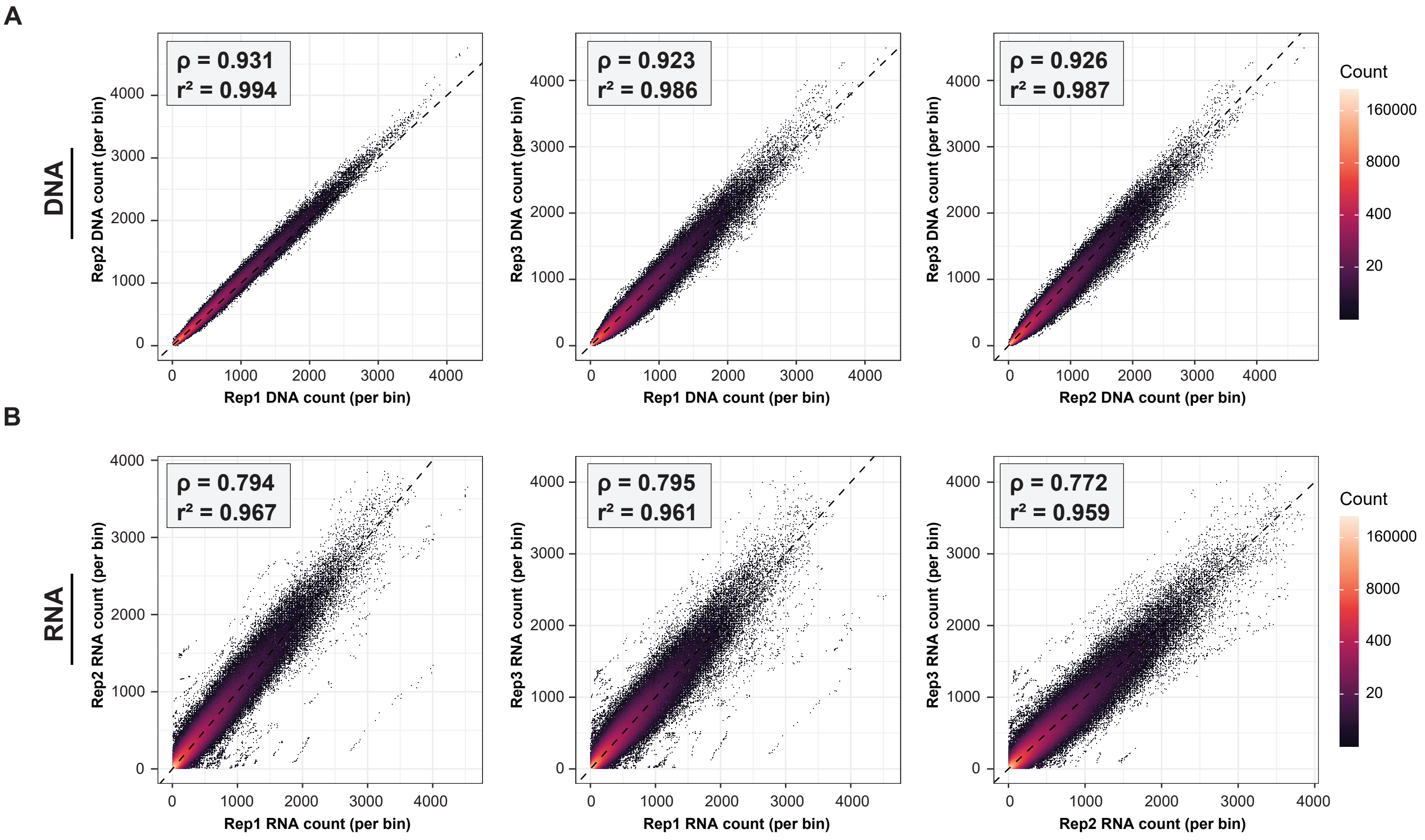


**Supplementary Figure 3. Coorelation between ATAC-STARR-seq replicates.** Scatter plots of DESeq2-normalized read counts per bin between replicates for both (A) DNA and (B) RNA samples. Pearson ($r^2$) and spearman ($\rho$) coorelation coefficients are indicated in the top left corner for each pairwise comparison.

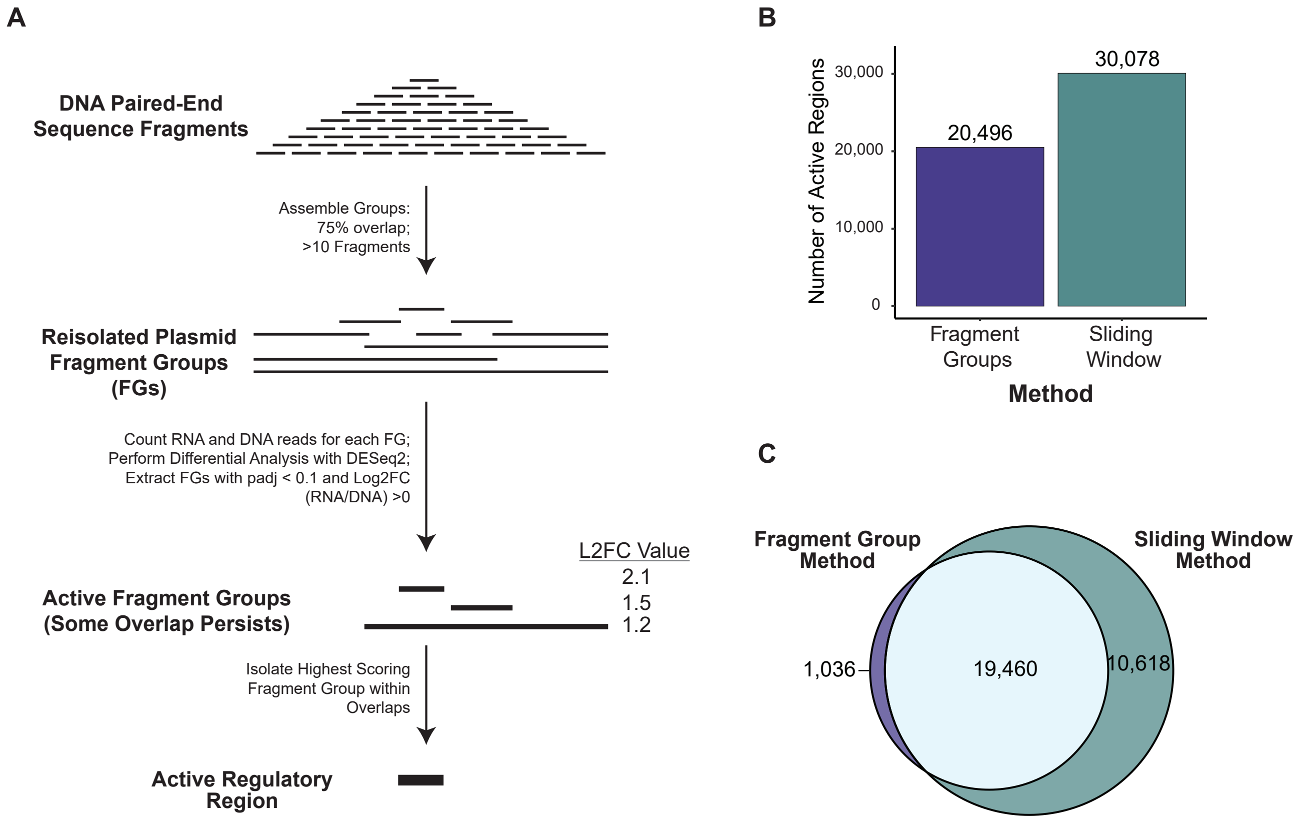


**Supplementary Figure 4. Comparison between the sliding window and the fragment group active region calling methods.** (A) Diagram of the fragment group region calling scheme. Paired-end fragments from the DNA samples are first assembled into "fragment groups" (FGs) which represent groups of more than 10 paired-end fragments with each fragment overlapping another fragment by at least 75%. Similar to the sliding window method, reads from RNA and DNA samples are then assigned to each FG and active FGs are identified using differential analysis with DESeq. The same padj (<0.05) and log2fold-change (>0) filters are applied. For FGs that overlap, the FG with the largest activity score is isolated. (B) The number of active regions called with either method. (C) Euler plot comparing the region overlap between the two methods.

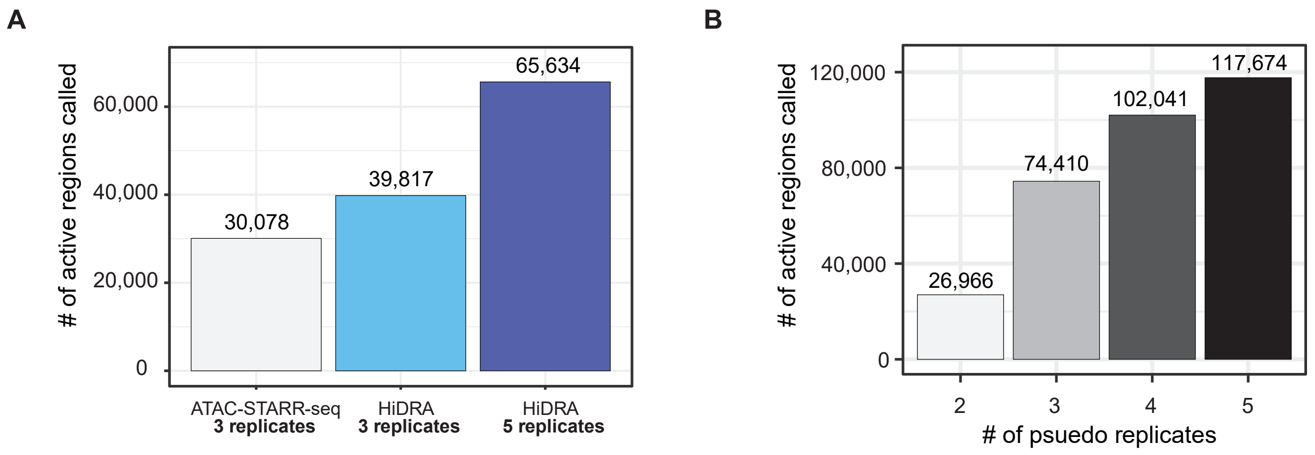


**Supplementary Figure 5. Analysis of replicate count on region calling sensitivity.** (A) Number of active regions called using HiDRA data with either 3 or 5 replicates. Our ATAC-STARR-seq active region number is plotted for comparison. (B) Number of active regions called when 2, 3, 4, or 5 pseudoreplicates are provided. To generate pseudoreplicates, replicates were merged and then split into 5 seperate files.

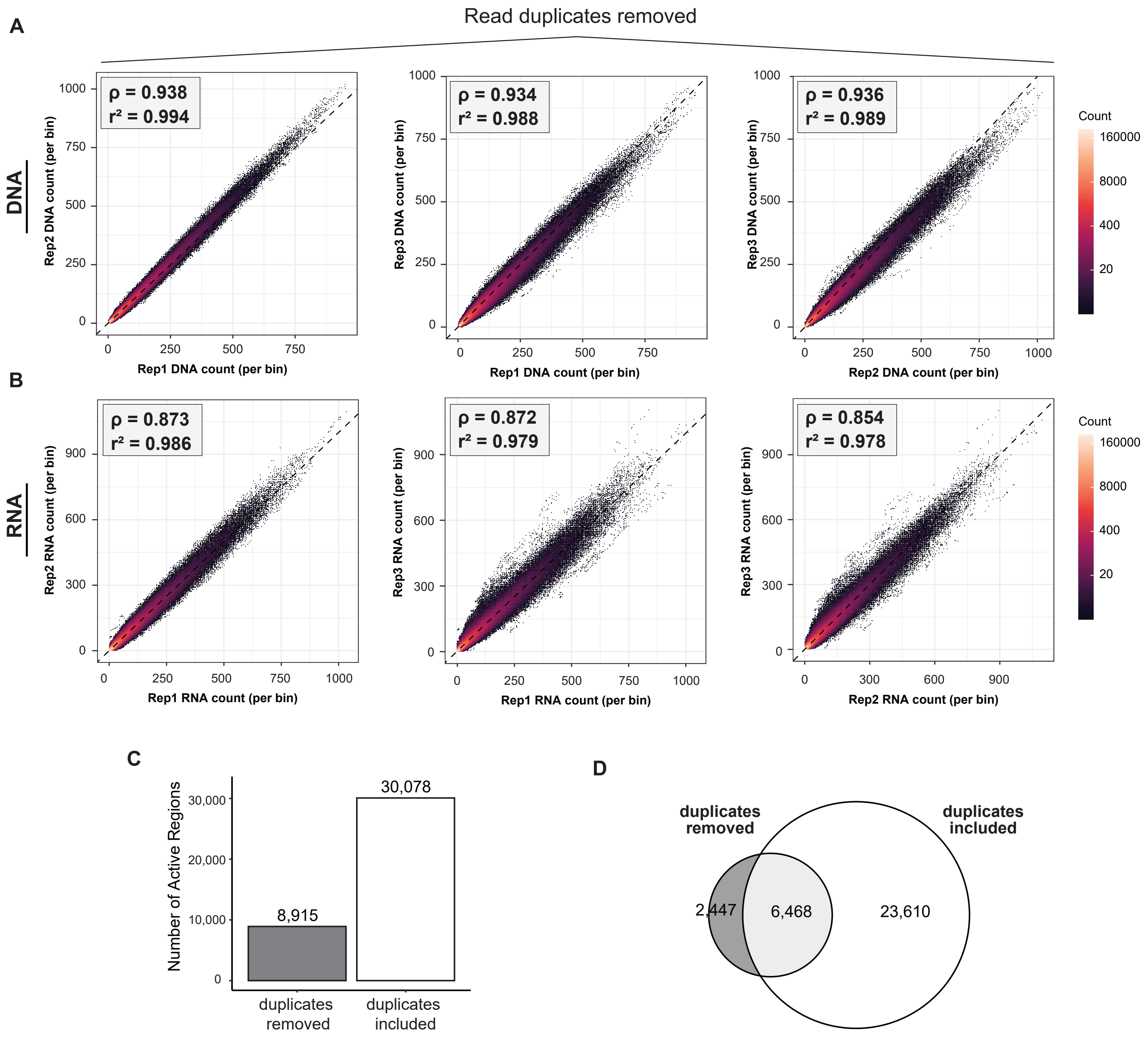


**Supplementary Figure 6. Comparison between keeping duplicates and removing duplicates to call active regions.** (A-B) Scatter plots of DESeq2-normalized read counts per bin between replicates for both (A) DNA and (B) RNA samples when duplicates are removed. Pearson ($r^2$) and spearman ($\rho$) coorelation coefficients are indicated in the top left corner for each pairwise comparison. (C) The number of active regions called with or without duplicates. (D) Euler plot comparing the region overlap between the two methods.

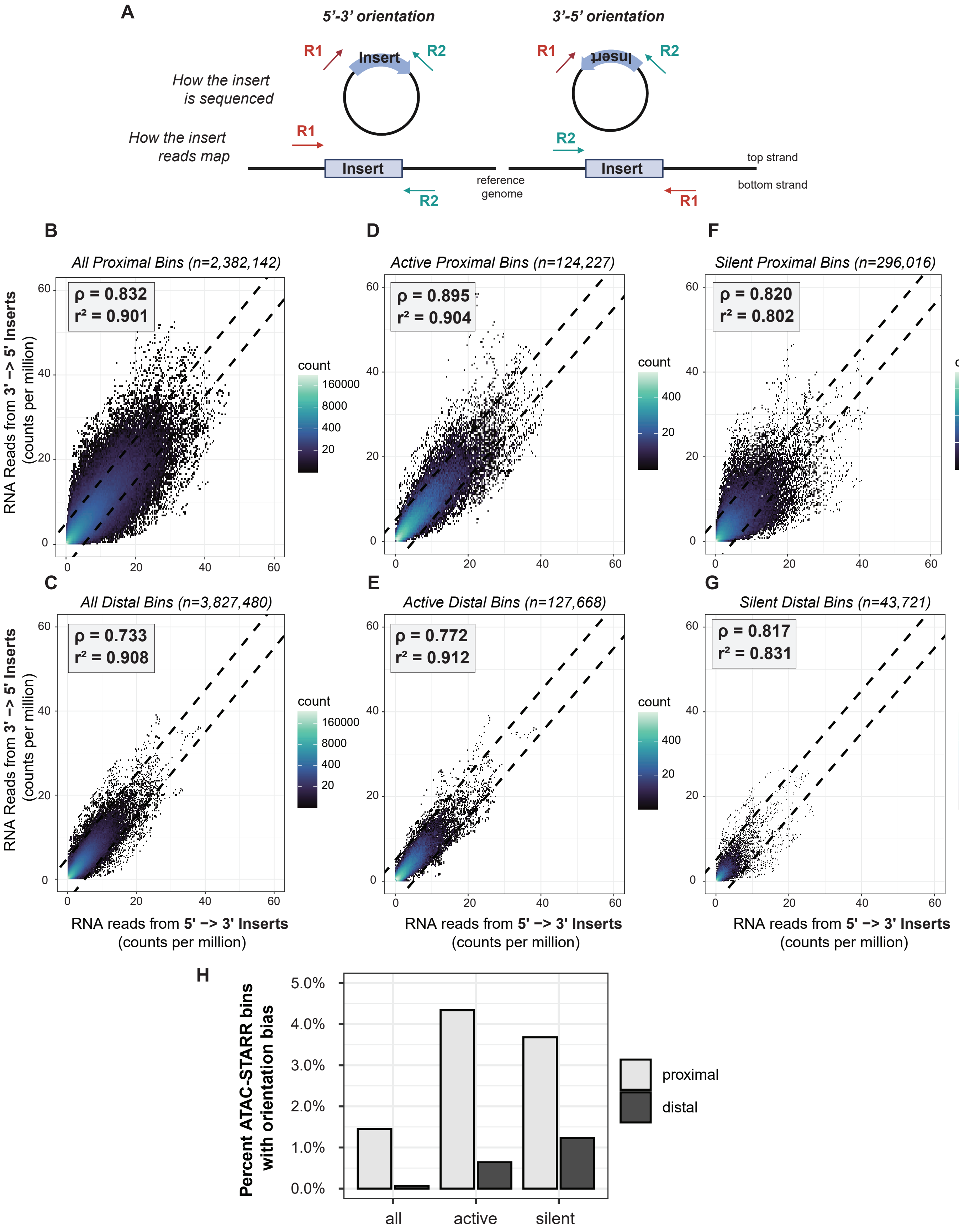


**Supplementary Figure 7. Assesement of potential orientation bias in ATAC-STARR-seq data.** (A) Schematic of the method for separating reads based on insert orientation. Read 1 and Read 2 are sequenced from the same position regardless of insert orientation on the plasmid and reporter RNA samples. Therefore, insert orientation can be specified based on how the read pair map to the genome. 5'-3' inserts have R1 on the top strand, while 3' -5' inserts have R1 on the bottom strand. (B-G) Scatter plots of counts per million normalized reporter RNA read counts between 5' to 3' inserts and 3' to 5' inserts for (B) all proximal bins analyzed, (C) all distal bins analyzed, (D) active proximal bins only, (E) active distal bins only, (F) silent proximal bins only or (G) silent distal bins only. Pearson ($r^2$) and spearman (ρ) coorelation coefficients are indicated in the top left corner for each pairwise comparison. Proximal bins were defined as within 2kb upstream and 1kb downstream of a transcription start site, while distal bins was defined as everything else. Dashed lines indicate +/- 5 counts from the expectation (y=x). The percentage of bins that lie outside of these lines are denoted in (H).

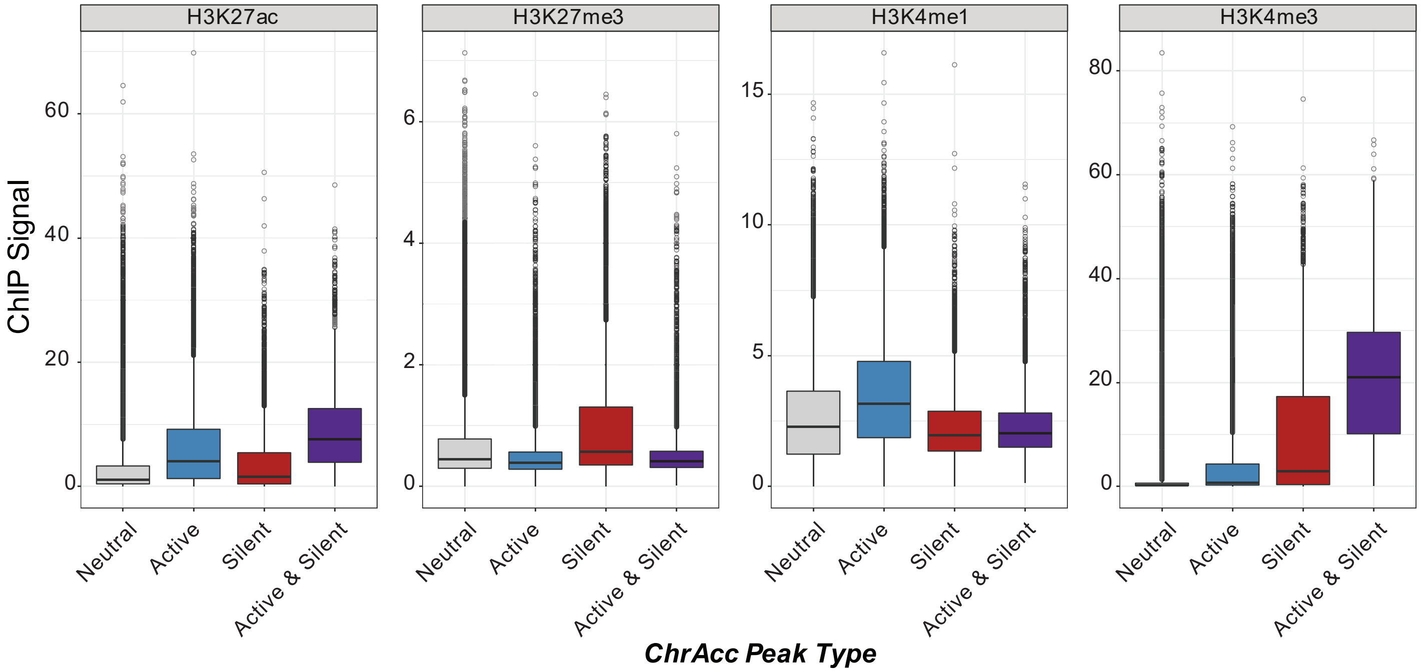


**Supplementary Figure 8. Histone modification ChIP-seq signal at accessible chromatin peaks.** Boxplot of the distribution of histone modification ChIP-seq signal for accessible chromatin peaks (ChrAcc) that contain an active region, a silent region, both and active and silent region, or neither (neutral). Values represents the average *fold change over control* signal per region for each histone modification.
